# Precision Enhancement of CAR-NK Cells through Non-Viral Engineering and Highly Multiplexed Base Editing

**DOI:** 10.1101/2024.03.05.582637

**Authors:** Minjing Wang, Joshua B. Krueger, Alexandria K. Gilkey, Erin M. Stelljes, Mitchell G. Kluesner, Emily J. Pomeroy, Joseph G. Skeate, Nicholas J. Slipek, Walker S. Lahr, Patricia N. Claudio Vázquez, Yueting Zhao, Ella J. Eaton, Kanut Laoharawee, Beau R. Webber, Branden S. Moriarity

## Abstract

Natural killer (NK) cells’ unique ability to kill transformed cells expressing stress ligands or lacking major histocompatibility complexes (MHC) has prompted their development for immunotherapy. However, NK cells have demonstrated only moderate responses against cancer in clinical trials and likely require advanced genome engineering to reach their full potential as a cancer therapeutic. Multiplex genome editing with CRISPR/Cas9 base editors (BE) has been used to enhance T cell function and has already entered clinical trials but has not been reported in human NK cells. Here, we report the first application of BE in primary NK cells to achieve both loss-of-function and gain-of-function mutations. We observed highly efficient single and multiplex base editing, resulting in significantly enhanced NK cell function. Next, we combined multiplex BE with non-viral *TcBuster* transposon-based integration to generate IL-15 armored CD19 CAR-NK cells with significantly improved functionality in a highly suppressive model of Burkitt’s lymphoma both *in vitro* and *in vivo*. The use of concomitant non-viral transposon engineering with multiplex base editing thus represents a highly versatile and efficient platform to generate CAR-NK products for cell-based immunotherapy and affords the flexibility to tailor multiple gene edits to maximize the effectiveness of the therapy for the cancer type being treated.

## Introduction

Natural killer (NK) cells have garnered extensive attention in immunotherapy due to their unique ability to kill transformed cells lacking major histocompatibility complexes (MHC) and antibody bound target cells through antigen-dependent cellular cytotoxicity (ADCC). From simply infusing allogeneic donor NK cells for treating hematological malignancies^1^ to adoptive therapy harnessing NK cells with chimeric antigen receptor (CAR) for a wide range of solid tumors, NK cells have exhibited many advantages over T cell-based therapy^2^. Despite promising early outcomes in clinical studies, rarely have NK cell therapies led to durable complete remission or tumor clearance^3,4^. Extensive studies have pointed to NK cell dysfunction, exhaustion, and lack of persistence as major contributing factors to the modest clinical outcomes^5^.

To date, engineering NK cells to express CAR constructs has relied almost exclusively on viral transduction with retroviruses^6^. Although highly efficient at delivering CAR constructs into NK cells, retrovirus cargo size limits and the risk of insertional oncogenesis present major drawbacks for clinical applications^6^. Comparatively, DNA transposon systems are less restricted by cargo size and afford a more economical and expeditious path to clinical manufacturing of CAR-NK cells^7^. Moreover, transposon systems allow for stable gene transfer in NK cells, which has distinct advantages over other non-viral systems, such as transient mRNA transfection^6^.

Limited *in vivo* persistence and durability issues in the absence of cytokine support are major challenges to effective NK-based immunotherapy^8^. Notably, NK cell persistence, metabolic fitness, and functionality have been enhanced through supplementation with exogenous IL-15 or IL-2 *in vitro* and *in vivo*^9–14^, suggesting gamma chain cytokine support may be a prerequisite to successful NK-based immunotherapies^8,15^. To this end, some NK-based clinical trials tested continuous intravenous infusion (CIV) of IL-15 to maintain NK cell persistence; however, this costly and toxic approach has driven researchers to develop IL-15 expressing NK cells, i.e. cytokine armored NK cells^15–18^. In fact, recent preclinical and clinical studies have demonstrated successful expression of soluble IL-15 (sIL-15) in CAR-NK cells using retroviral engineering, which led to improved cytotoxicity and persistence *in vivo*^17–19^.

Beyond cytokine armoring, we^20^ and others^21–24^ have demonstrated that CRISPR/Cas9 can be utilized to enhance NK cell function through targeted gene knockout (KO). However, this approach is not ideal for multiplex gene KO as translocations and other genomic rearrangements can occur due to simultaneous induction of multiple double strand breaks (DSBs). Due to these concerns, we previously deployed base editor technology in primary human T cells to enhance CAR-T cell function through multiplex gene KO^25^. Base editors consist of a catalytically inactive Cas9 fused to a deaminase domain for site specific nucleotide base conversion, allowing users to install gain or loss of function mutations without induction of a DSB or requirement for a DNA donor molecule^26–29^. However, BEs have yet to be deployed for primary NK cell engineering to achieve similar multiplex base editing.

Here, we report the first highly efficient engineering of NK cells using an adenine base editor (ABE), ABE8e^30^, to achieve both loss-of-function and gain-of-function mutations in NK cells to improve their function. We also explored the upper limit of simultaneous multiplex base editing and observed no significant reduction in base editing efficiency targeting up to 6 independent loci. Next, we identified optimal, synergistic multiplex KO combinations and co-delivered a CD19 CAR and IL-15 armoring using the *TcBuster* DNA transposon system to develop a non-viral engineered CAR-NK cell product tailored specifically to overcome a highly suppressive model of Burkitt’s lymphoma. Our work serves as a proof of concept that precision enhancement of CAR-NK cells can be achieved using simultaneous multiplex base editing and non-viral transposon-based engineering to generate CAR-NK therapies tailored for user defined cancer types.

## Results

### Highly efficient single gene KO in primary human NK cells using ABE

Base editors (BEs) are capable of highly efficient gene KO without double-stranded break induction through splice site disruption or installations of stop codons^25,31–35^. We first examined whether single gene KO could be achieved using BE in primary NK cells and evaluated consequent changes in immune function. Given that suppression of inhibitory signals is one of the most common strategies for enhancing NK-based therapy^36^, we assembled a panel of targets focusing largely on intracellular and extracellular inhibitory proteins expressed by NK cells **(Table 1)**. Intracellular checkpoints include AHR, a negative regulator of NK cell cytotoxicity when agonists (i.e. kynurenine) are present in the tumor microenvironment (TME)^37^, and cytokine-inducible SH2-containing protein (CISH), a negative regulator of IL-15 signaling essential for NK cell development and function^34^. On the surface of NK cells, programmed cell death protein 1 (*PDCD1*) and T-cell immunoreceptor with Ig and ITIM domains (*TIGIT*) are well-defined inhibitory immune checkpoints whose signal blockade has been seen a wide range of clinical applications^38,39^. KLRG1, an emerging target in immunotherapy, is an additional inhibitory immune checkpoint with potential clinical relevance^40^. Since our previous BE studies found targeting splice-donors (SD) is the most efficient method for gene disruption, we designed sgRNAs targeting a SD in each gene of interest using SpliceR, an in-house-developed online BE sgRNA design and prediction tool^35^. Each sgRNA was co-transfected with ABE8e mRNA into feeder-stimulated peripheral blood (PB) NK cells isolated from healthy donors by electroporation. KO efficiency was then assessed at the genomic level through Sanger sequencing and quantification of A to G conversion rates using EditR software **(Figure 1a)**^41^. Specifically, A to G conversion rates for each targeted gene were as follows: *AHR* 100% (± 0%), *CISH* 99.67% (± 0.58%), *KLRG1* 65.07% (± 14.86%), *TIGIT* 100% (± 0%), *PDCD1* 86.33% (± 7.51%). Example sequencing results of each target gene as assessed by EditR are shown in **Supplement Figure 1a-e**^41^. Of note, indels were not detectable for any of the target genes in all 3 donors **(Supplement Figure 2)**. Next, we assessed protein level KO of each target gene via western blot (WB) or flow cytometry using donor-matched control NK cells **(Figure 1b & Supplement Figure 3a-d)**. For intracellular proteins, WB analysis showed an average protein reduction of 95.60% (± 3.88%) for AhR, and 99.59% (± 0.71%) for CIS. Flow analysis demonstrated a 96.78% (± 4.41%) reduction of TIGIT surface expression, while KLRG1 showed an average of 65% (± 14.86%) reduction in protein expression. Note that PD-1 protein loss results are not shown here, as minimal detection of PD-1 was achieved in all healthy donors used. This observation is in line with previous studies reporting low *detectable* expression (<5%) of PD-1 in NK cells of healthy individuals and elevated expression in patients bearing tumors^42–50^.

**Fig. 1.**
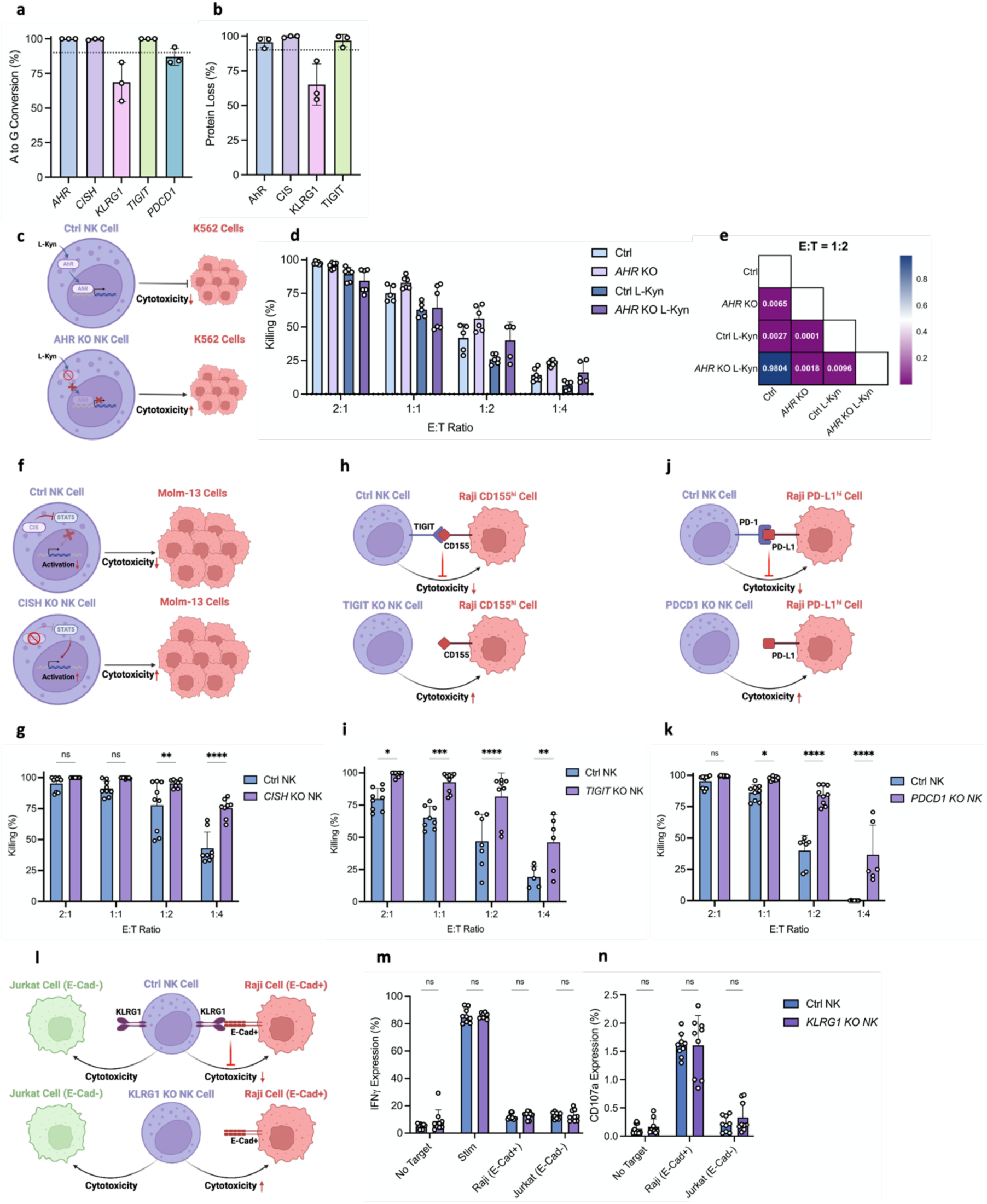
Highly efficient single gene KO in NK cells using BE. **a** Editing efficiency at genomic level quantified by A to G conversion of target base for each gene locus (n=3 independent NK cell donors). **b** Editing efficiency at protein level quantified by percentage of protein loss of each gene (n=3 independent NK cell donors). **c** Schema of killing assay to assess the functional improvement of AHR KO NK cells. **d** Ability of AHR KO versus Ctrl NK cells (with or without 14 days of L- Kynurenine pre-treatment) to kill K562 cells at various E:T ratios as measured by luciferase luminescence assay. Assays run in triplicate in n=2 independent biological NK cell donors. **e** Statistical significance (P-value) between each condition of AHR KO functional killing assay at E to T ratio of 1:2. **f** Schema of killing assay to assess the functional improvement of CISH KO NK cells. **g** Ability of CISH KO versus Ctrl NK cells to kill Molm-13 cells at various E:T ratios as measured by luciferase luminescence assay. Assays run in triplicate in n=3 independent biological NK cell donors. **h** Schema of killing assay to assess the functional improvement of TIGIT KO NK cells. **i** Ability of TIGIT KO versus Ctrl NK cells to kill Raji CD155^hi^ cells at various E:T ratios as measured by luciferase luminescence assay. Assays run in triplicate in n=3 independent biological NK cell donors. **j** Schema of killing assay to assess the functional improvement of PDCD1 KO NK cells. **k** Ability of PDCD1 KO versus Ctrl NK cells to kill Raji PD-L1^hi^ cells at various E:T ratios as measured by luciferase luminescence assay. Assays run in triplicate in n=3 independent biological NK cell donors. **l** Schema of ICS assay to assess the functional improvement of KLRG1 KO NK cells. **m&n** Cytokine production (**m**) and degranulation (**n**) ability of KLRG1 KO versus Ctrl NK cells against E-Cad+ Raji cells or E-Cad- Jurkat cells as measured by percentage of NK cells produce IFN𝛾 and CD107a. Assays run in triplicate in n=3 independent biological NK cell donors. Data represented as mean ± SD. P-values calculated by two-way ANOVA test (n.s.P > 0.05, *P ≤ 0.05, **P ≤ 0.01, ***P ≤ 0.001, ****P ≤ 0.0001).

**Table 1.**
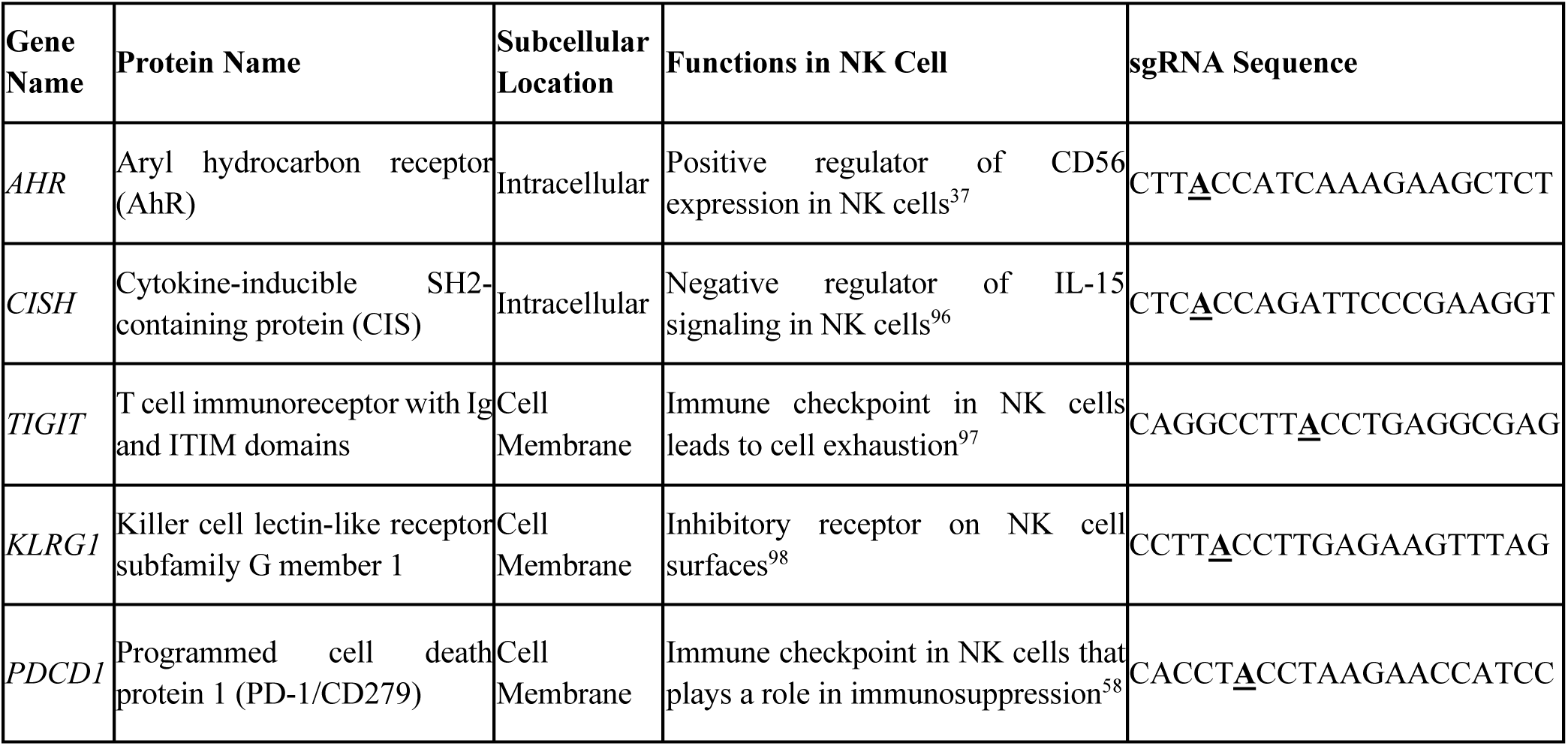
List of genes for KO in NK cells using BE.

We next assessed whether individual gene KOs translated into the expected functional enhancement of NK cells by designing functional assays specific for each gene target. Starting with the intracellular targets, AHR agonists are known to suppress NK cell cytotoxicity during tumor progression and metastasis^37,51^. Thus, we cultured NK cells with an AHR agonist, L-Kynurenine (L-Kyn), for two rounds of 7-day expansion (14 days) before co-culturing them with K562 target cells at various effector to target (E:T) ratios **(Figure 1c)**. Our results showed that L-Kyn significantly reduced NK cell killing overall, however, *AHR* KO NK exhibited significantly higher levels of killing than control NK **(Figure 1d & e and Supplement Figure 4a-d)**. Due to its role in negative regulation of IL-15 signaling, several groups reported improved functionality of NK cells with *CISH* KO^34,52,53^. Here, we co-cultured *CISH* KO and control NK cells with an aggressive acute myeloid leukemia (AML) cell line, Molm-13, for 48 hours at multiple E:T ratios **(Figure 1f)**. Compared to control NK, *CISH* KO NK exhibited enhanced cytotoxicity against Molm-13 cells, in line with previous reports **(Figure 1g)**^34,52,53^.

Next, we examined the functional impacts of surface checkpoint KOs. TIGIT/CD155 interaction serves as an inhibitory immune checkpoint in NK cells, and other studies have demonstrated enhanced NK cell cytotoxicity with *TIGIT* KO^54–56^. In line with these reports, BE mediated *TIGIT* KO NK cells co-cultured with Raji cells engineered to overexpress CD155 (i.e. Raji CD155^hi^ cells) for 48 hours at different E:T ratios exhibited significantly higher cytotoxicity against Raji CD155^hi^ cells compared to donor-matched control NK cells **(Figure 1h, i)**. Analogous to TIGIT/CD155, the PD-1/PD-L1 axis is another well-known inhibitory immune checkpoint in NK cells^57–60^. BE mediated *PDCD1* KO NK cells co-cultured with Raji cells engineered to overexpress express PD- L1 (i.e. PD-L1^hi^ cells) showed enhanced killing versus control NK cells **(Figure 1j, k)**. Finally, E-cadherin (E-Cad) binding to KLRG1 has been shown to protect tumor cells from cytolysis by NK cells^61^. Thus, we co-cultured NK cells with either E-Cad(+) cells (Raji) or E-Cad(-) cells (Jurkat) and performed intracellular cytokine staining (ICS) to assess activation **(Figure 1l)**. In contrast to previous reports, we observed no significant differences in CD107a positivity and interferon-gamma (IFN-γ) expression between control NK and KLRG1 KO NK against E-Cad(+) or E-Cad(-) target cells **(Figure 1m, n)**. In summary, using ABE8e we were able to achieve highly efficient single gene KO in NK cells, which translated into *in vitro* functional enhancement against multiple tumor models, with the exception of *KLRG1* KO.

### Highly efficient installation of a non-cleavable CD16A mutation in NK cells using ABE

Given that BE can target individual nucleotides to alter amino acid codons, we next sought to establish whether ABE could be used to enhance NK cell function through gain-of-function mutations. CD16a is an Fc receptor expressed on the surface of NK cells which mediates antibody-dependent cell-mediated cytotoxicity (ADCC)^62^. Upon NK cell activation, CD16a undergoes a proteolytic cleavage process by ADAM17 as a negative feedback mechanism regulating NK cell cytotoxicity **(Figure 2a)**^63^. Previous studies have found that a single amino acid substitution (S197P) at the cleavage site of ADAM17 on CD16a rendered the receptor “non-cleavable”, and expression of the non-cleavable CD16a (ncCD16a) cDNA in NK cells enhanced ADCC^64^. Here, we sought to install the non-cleavable CD16a (ncCD16a) mutation at the endogenous locus using ABE8e as a proof-of-principle of ABE-based gain-of-function engineering in NK cells. To this end, we designed a sgRNA targeting serine197 at the ADAM17 cleavage site with the goal of replicating the S197P mutation through a single A to G base conversion, altering the mRNA codon from UCA to CCA **(Figure 2b)**. Sequencing results validated our prediction and yielded an average A to G conversion rate of 89% (± 9.17%, n=3) in NK cells with the highest conversion rate observed at 99% in one donor **(Figure 2c; Supplement Figure 5a)**. To assess whether the installation of this mutation conferred cleavage resistance, NK cells were stimulated with phorbol 12-myristate 13-acetate (PMA) to activate ADAM17 and then assessed for CD16a retention (i.e. % of CD16a detected with PMA treatment versus % of CD16a detected without PMA treatment). As shown in **Figure 2d & Supplement Figure 5b**, we observed an average of 97.17% (± 2.45%) CD16a retention after PMA treatment in ABE8e engineered NK cells, with the highest retention rate approaching 100%.

**Fig. 2.**
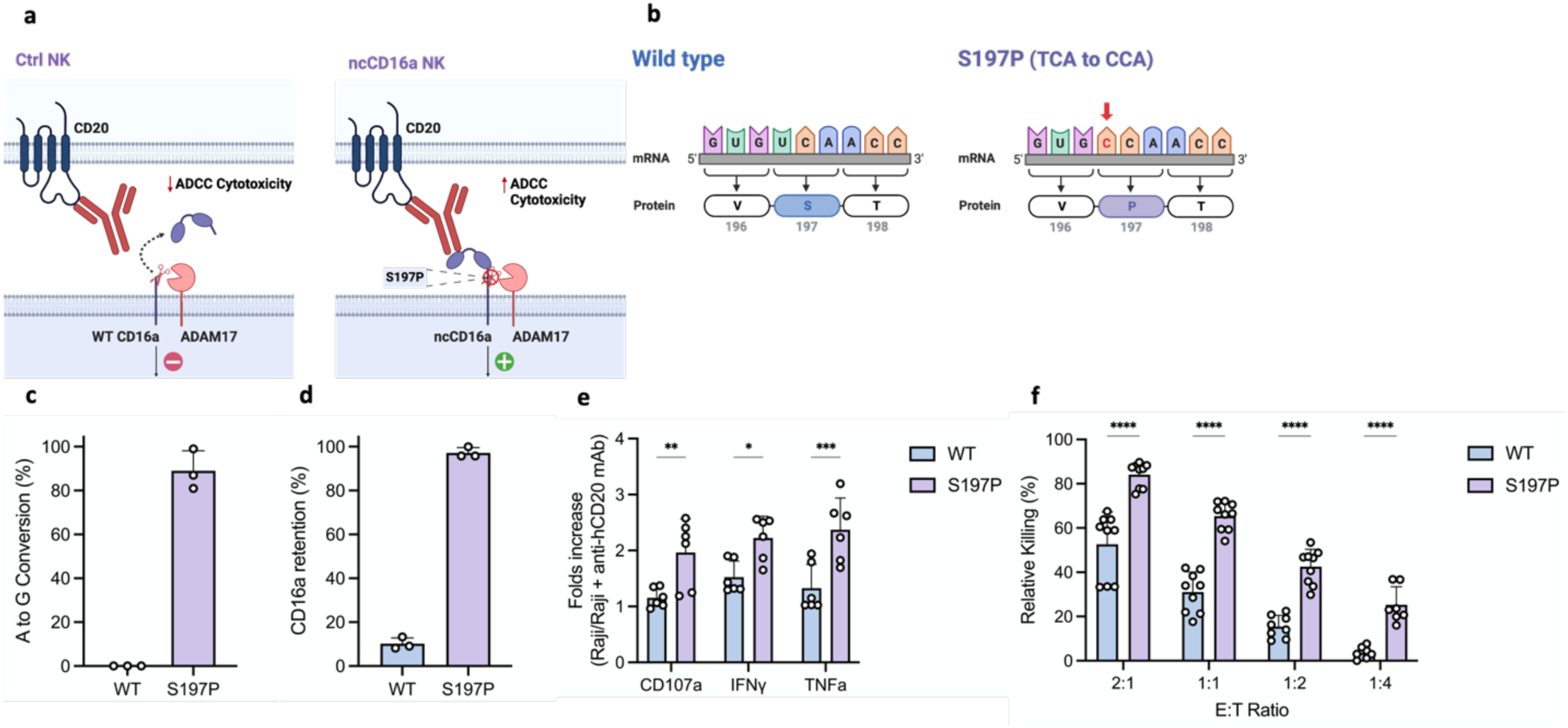
Highly efficient gain-of-function mutation in NK cells using BE. **a** Schema of how the S197P mutation renders NK cells non-cleavable by ADAM17 and results in enhanced ADCC cytotoxicity. **b** Schema of the recreation of S197P ncCD16a NK cells by a single base modification by BE. **c** Editing efficiency at genomic level quantified by A to G conversion of target base for *CD16A* (n=3 independent NK cell donors). **d** Editing efficiency at protein level quantified by CD16a retention on NK cell surface after PMA treatment (n=3 independent NK cell donors). **e** Cytokine production of S197P CD16a versus WT CD16a NK cells against CD20+ Raji cells during ADCC (E:T ratio: 1:1). Plotted by folds increase of each cytokine with versus without anti- hCD20 mAb co-treatment. Assays run in duplicate in n=3 independent biological NK cell donors. **f** Ability of S197P CD16a versus WT CD16a NK cells to carry out ADCC against anti-hCD20 mAb treated CD20+ Raji cells at various E:T ratios as measured by luciferase luminescence assay. Assays run in triplicate in n=3 independent biological NK cell donors. Data represented as mean ± SD. P-values calculated by two-way ANOVA test (*P ≤ 0.05, **P ≤ 0.01, ***P ≤ 0.001, ****P ≤ 0.0001)

Next, to validate the functionality of our ncCD16a NK cells, we tested their ability to mediate ADCC against Raji cells treated with an anti-hCD20-mAb. We first assessed activation of ncCD16a NK cells in this assay via ICS for degranulation (CD107a) and cytokine production (TNF-α and IFN-γ). Compared to non-engineered NK cells, ncCD16a NK cells demonstrated significantly increased CD107a, IFN-γ, and TNF-α expression when co-cultured with cell targets treated with anti-hCD20-mAb, indicating enhanced ADCC-driven activation (measured by Raji versus Raji co-treatment with anti-hCD20-mAb) of ncCD16a NK cells **(Figure 2e)**. Functionally, we assessed NK cytotoxicity via ADCC using an ADCC co-culture killing assay against Raji cells at different E:T ratios. As predicted, ncCD16a NK exhibited significantly enhanced killing of Raji cells compared to control NK cells throughout all E:T ratios tested **(Figure 2f)**. Surprisingly, when co-cultured without anti-hCD20-mAb, ncCD16a NK also demonstrated significantly enhanced killing compared to that of control NK cells **(Supplement Figure 5c)**. Taken together, these data demonstrate that ABE is capable of installing gain-of-function mutations in NK cells at high efficiencies to improve NK cell functionality.

### Highly efficient multiplex base editing in NK cells using ABE

Next generation cellular therapies will likely require a multitude of tailored gene edits to maximize efficiency, particularly when addressing hard to treat solid tumor cancers. Previously, our group has shown multiplex base editing can be used to enhance CAR-T cell function^25^. Here, we examined whether the same approach can be applied in developing NK-based cell therapies using ABE8e and also attempted to find the upper limits of multiplex base editing. We started with one sgRNA targeting *TIGIT* and subsequently added sgRNAs one at a time in the following order: ncCD16a sgRNA, AHR sgRNA, KLRG1 sgRNA, CISH sgRNA, PDCD1 sgRNA up to a total of 6 sgRNAs (**Figure 3a)**. Multiplex editing efficiencies at the genomic level were again assessed via Sanger sequencing of A to G conversions at target sites assessed using EditR. Impressively, we observed no significant loss of editing efficiency at any loci using up to 6 sgRNAs simultaneously **(Figure 3b and Supplement Figure 6a-f)**. Editing efficiencies at protein level, measured by protein loss (for AhR, CISH, KLRG1, TIGIT, and PD-1) or CD16a retention rate after PMA treatment were consistent with genomic base editing levels **(Figure 3c and Supplement Figure 7a-e)**. In summary, these data demonstrate that highly efficient multiplex base editing is achievable in NK cells using ABE and the upper limit of simultaneous base edits has not yet been reached in this context.

**Fig. 3.**
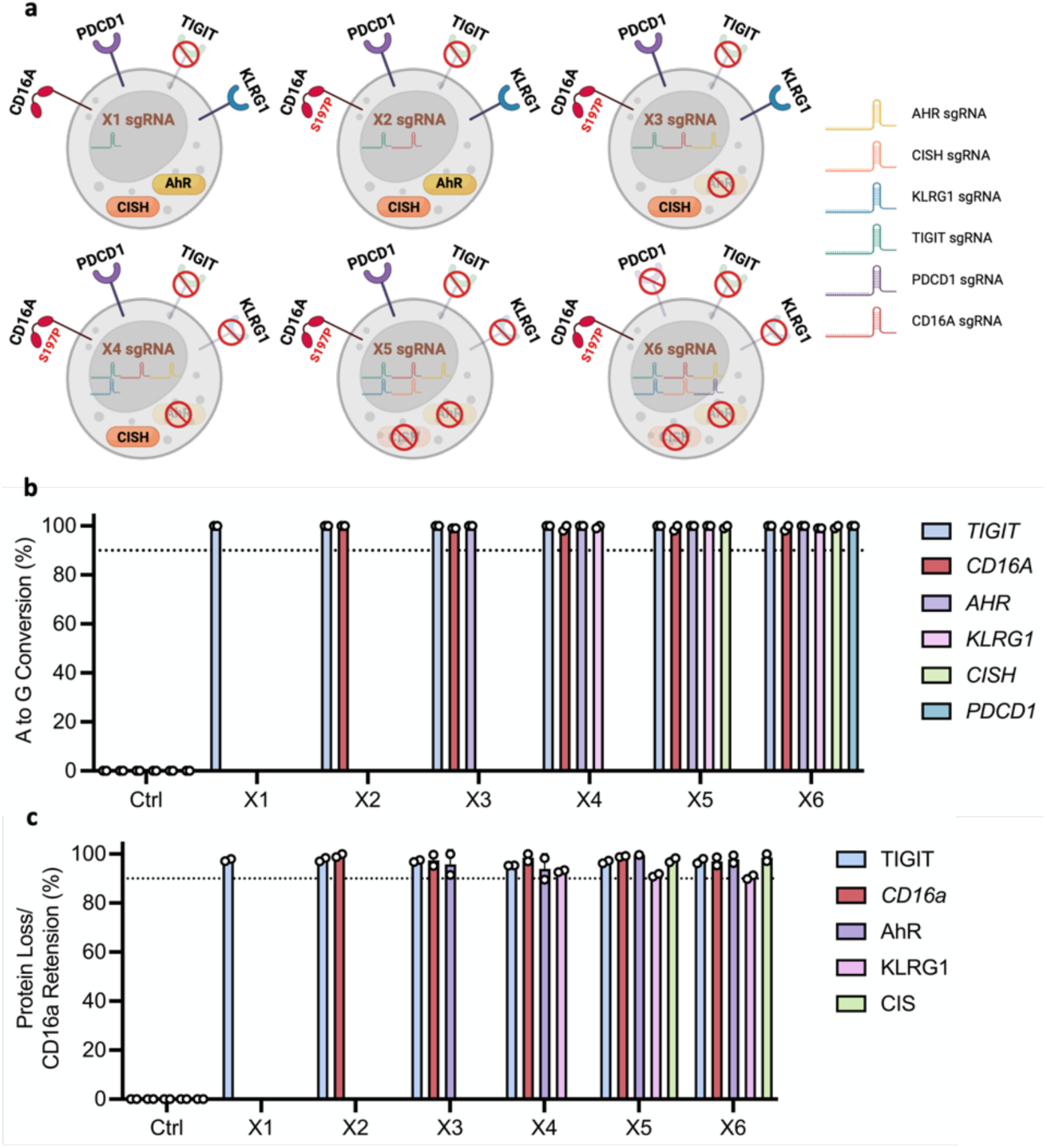
Highly efficient multiplex editing in NK cells using BE. **a** Schema of multiplex editing strategy. **b** Multiplex editing efficiency at genomic level quantified by A to G conversion of target base for each gene locus (n=2 independent NK cell donors). **c** Multiplex editing efficiency at protein level quantified by percentage of protein loss of each gene (n=2 independent NK cell donors). Data represented as mean ± SD.

### Tailored multiplex engineering of NK cells for optimized cytotoxicity against Raji CD155^hi^/PD-L1^hi^ cells

In order to demonstrate a tailored editing approach for NK based cancer immunotherapy, we next sought to determine an optimal combination of multiplex edits that yields the most potent NK cell cytotoxicity for a chosen tumor model. Two of the most common factors leading to poor prognosis in many cancers are high expression of PD-L1 and CD155^65–73^. To model this, we generated a Raji cell line that over-expresses both checkpoint ligands (i.e. Raji^hi/hi^) **(Supplement Figure 8)**. To determine the optimal multiplex base editing combination, we started with TIGIT (CD155 ligand) and PDCD1 (PD-L1 ligand) KO to avoid suppression from CD155 and PD1, respectively. Next, these edits were combined with AHR KO, CISH KO, or both KOs for testing different base editing combinations **(Figure 4a)**. Editing efficiencies showed the average A to G conversions of each target site in all combinations were above 90%, and similar results were observed for average protein loss of all KOs **(Figure 4b, c)**. In order to assess the functional consequences of the various combinations of base edits, we first performed co-culture killing assays with Raji^hi/hi^ cells at different E:T ratios. We observed that unedited NK cells had the lowest level of killing against Raji^hi/hi^ and there was no significant difference in killing between TIGIT+PDCD1 (TP) KO NK, TIGIT+PDCD1+AHR (TPA) KO NK, and TIGIT+PDCD1+AHR+CISH (TPAC) KO NK. However, knocking out TIGIT, PDCD1, and CISH (TPC) demonstrated the highest killing against Raji^hi/hi^ cells compared to all other combinations **(Figure 4d-e and Supplement Figure 9a-d)**. These data demonstrate that TPC KO NK cells have the most robust cytotoxicity against Raji^hi/hi^ cells compared to all other multiplex base editing combinations.

**Fig. 4.**
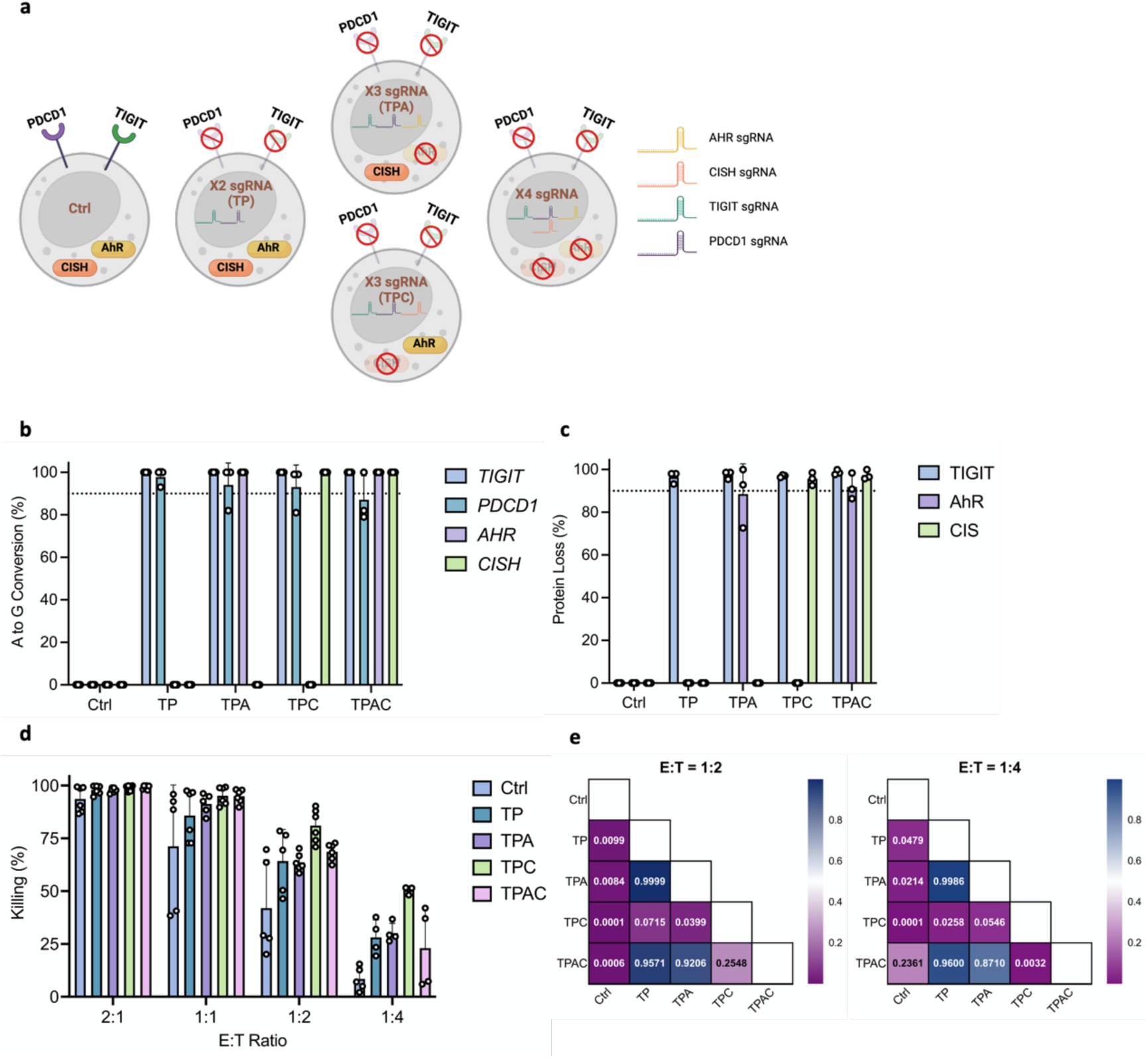
Optimization of multiplex KO to maximize NK functionality against Raji^hi/hi^. **a** Schema of optimization strategy and all possible KO combinations included. **b** Editing efficiency at genomic level quantified by A to G conversion of target base for each gene locus (n=3 independent NK cell donors). **c** Editing efficiency at protein level quantified by percentage of protein loss of each gene (n=3 independent NK cell donors). **d** Ability of each KO combination to kill Raji^hi/hi^ cells at various E:T ratios as measured by luciferase luminescence assay. Assays run in triplicate in n=2 independent biological NK cell donors. **e** Functional killing assay statistical significance (P-value) between each KO combination at E to T ratio of 1:2 and 1:4. Ctrl: ABE8e mRNA only, TP: *TIGIT* and *PDCD1* KO, TPA: *TIGIT*, *PDCD1*, and *AHR* KO, TPC: *TIGIT*, *PDCD1*, and *CISH* KO, TPAC: *TIGIT*, *PDCD1*, *AHR,* and *CISH* KO. Data represented as mean ± SD. P-values calculated by two-way ANOVA test.

### Simultaneous multiplex base editing and non-viral transposon engineering generates highly functional CAR-NK cells

Chimeric antigen receptors (CARs) have emerged as one of the most promising tools for enhancing cell-based cancer immunotherapy in the past decade^74^. To date, delivery of CAR constructs into NK cells has been largely limited to the use of viral vectors^75–77^. The cost and extended manufacturing process of viral vectors drives the needs for a faster and more economical alternative method for stable transgene delivery. Thus, to develop an effective non-viral engineered CAR-NK cell product, we designed a *TcBuster* transposon plasmid encoding a CD19 CAR **(Figure 5a; ‘CAR’).** In an effort to test whether cytokine armoring of CAR-NKs would enhance persistence and activity, we designed an additional transposon plasmid encoding sIL15 **(Figure 5a; ‘CAR15’)**. Notably, IL15 has been shown to be necessary to support NK cell persistence in NOD-scid IL2R gamma-null (NSG) models^78,79^. Each transposon contains the MND promoter, 2^nd^ generation CD19-targeting CAR, and RQR8 **(Figure 5a)**. RQR8 is a synthetic cell surface receptor with CD34 and CD20 epitopes and was included here for flow cytometry detection and immunomagnetic enrichment of CAR-expressing NK cells^80^.

**Fig. 5.**
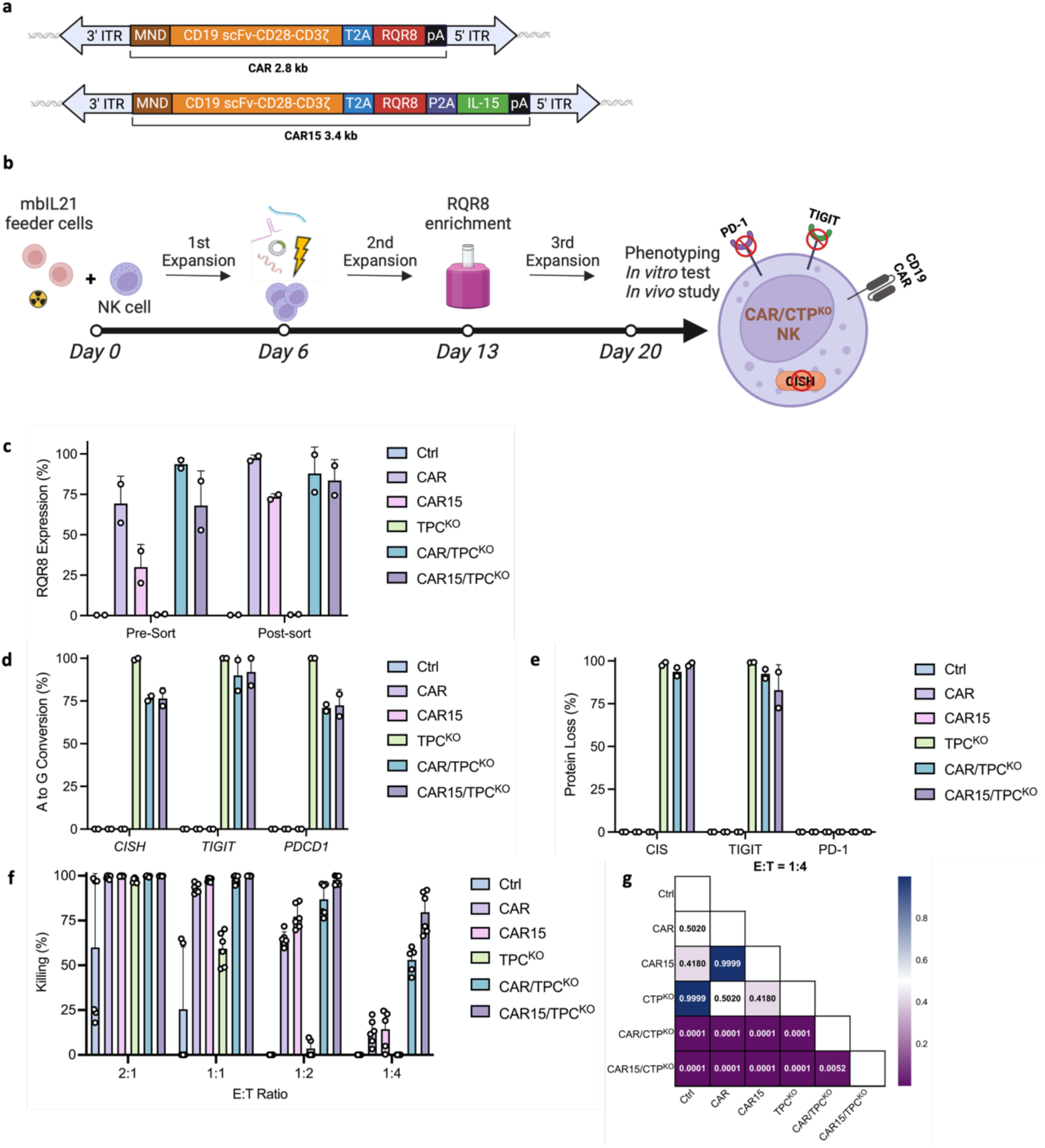
Simultaneous BE and non-viral transposon engineering exhibited enhanced NK cytotoxicity. **a** Schema of the designs of CD19 CAR constructs. **b** Schema of NK cell engineering timeline for simultaneous sgRNAs and CD19 CAR delivery. **c** Pre-sorting and post-sorting CAR integration rate quantified by percentage of RQR8 expression on NK cells (n=2 independent NK cell donors). **d** Post-sorting editing efficiency at genomic level quantified by A to G conversion of target base for each gene locus (n=2 independent NK cell donors). **e** Post-sorting editing efficiency at protein level quantified by percentage of protein loss of each gene (n=2 independent NK cell donors). **f** *In vitro* testing of the cytotoxicity of simultaneous BE and TcBuster engineered NK cells against Raji^hi/hi^ cells at various E:T ratios as measured by luciferase luminescence assay. Assays run in triplicate in n=2 independent biological NK cell donors. **g** Killing assay statistical significance (P-value) between each condition at E to T ratio of 1:4. Ctrl: ABE8e mRNA and CAR-expressing nanoplasmid only, CAR: CD19 CAR RQR8; CAR15: CD19 CAR RQR8 IL-15; CTP^KO^: *CISH*, *TIGIT*, and *PDCD1* KO; CAR/CTP^KO^: CD19 CAR RQR8 with *CISH*, *TIGIT*, and *PDCD1* KO; CAR15/CTP^KO^: CD19 CAR RQR8 IL-15 with *CISH*, *TIGIT*, and *PDCD1* KO. Data represented as mean ± SD. P-values calculated by two-way ANOVA test.

For engineering, NK cells were feeder expanded before co-delivery of CD19 CAR transposon plasmid, *TcBuster* transposase mRNA, sgRNAs, and ABE8e mRNA through electroporation **(Figure 5b)**. Following a subsequent 7-day NK cell feeder expansion, CD19 CAR integration rate was assessed via flow cytometry before CAR enrichment through RQR8 immunomagnetic selection **(Figure 5c)**. CAR integration rates in each group were reasonably efficient in multiple donors (57.4%-81.3% for CAR and 20.1%-39.9% for CAR15) and comparable to previously reported integration rates of CD19 CAR through retroviral transduction^75,77^. We also assessed the insertion site profile of *TcBuster* transposons and found that it integrated in coding genes less frequently than lentivirus or retrovirus regardless of construct **(Supplement Figure 10a)**. Additionally, evaluation of the distance between integration events and transcriptional start sites indicated that *TcBuster* integrated further away from transcriptional start sites when compared to lentivirus and retrovirus **(Supplement Figure 10b)**. CAR enrichment was further performed to ensure CAR-expression across groups was not significantly different, which could confound results comparing across groups. After a third and final feeder-based expansion, CD19 CAR expression was assessed by flow cytometry **(Figure 5c)** and IL-15 expression was confirmed through ELISA **(Supplement Figure 11;** CAR15: 220.0 ± 5.09 pg/mL, CAR15/TPC^KO^: 340.9 ± 20.34 pg/mL). Control groups lacking transposon engineering demonstrated base editing levels comparable to previous results in TPC KO NK cells **(Figure 4b)**, while a 10-25% reduction in base editing efficiency was observed in groups with transposon co-delivery **(Figure 5d)**. However, the average protein reduction of each gene of interest in all KO groups was above 90%, with one outlier NK donor from the CAR15/TPC^KO^ group having lower TIGIT protein reduction **(Figure 5e)**.

The functionality of all engineered NK cell groups was first assessed using co-culture killing assays with Raji^hi/hi^ cells at a range of E:T ratios. We observed no significant differences in killing between CAR and CAR15 groups, but a moderate enhancement of killing by CAR15/TPC^KO^ group compared to CAR/TPC^KO^ NK cells **(Figure 5f-g and Supplement Figure 12a-d)**. TPC^KO^ also improved cytotoxicity of NK cells, with both CAR/TPC^KO^ and CAR15/TPC^KO^ groups exhibiting better killing against Raji^hi/hi^ cells when compared to that of their corresponding CD19 CAR only groups. These data demonstrate that CAR15/TPC^KO^ NK cells exhibited the most significant functionality improvement against Raji^hi/hi^ targets *in vitro*.

### Multiplex base edited CD19 CAR-NK cells are highly functional *in vivo*

To test the efficacy of our multiplex base edited CAR-NK cell product *in vivo*, we used Raji^hi/hi^ cells as a highly suppressive xenograft model of Burkitt’s lymphoma. NSG mice were xenografted with 1E5 of Raji^hi/hi^ through intravenous injection (I.V.) on day 0, followed by a randomization into treatment groups of equivalent tumor burden prior to therapy delivery **(Supplement Figure 13a)**. Mice receiving therapy were dosed with two rounds of engineered NK cells 14 days apart (day 3 and 17) **(Figure 6a)**. Mouse body weight and bioluminescent imaging (BLI) of tumor burden were monitored weekly until predefined humane endpoints were reached **(Figure 6b and Supplement Figure 13b)**. Bioluminescent imaging results suggested tumor progression was significantly delayed in groups treated with CAR15 NK (n=5) and CAR15/TPC^KO^ NK (n=5) compared to all other groups **(Figure 6b, c)**. Mouse tumor burden of these two groups was significantly reduced by day 23, the last imaging time point before mice began to meet endpoint criteria **(Figure 6d)**. The survival of mice receiving CAR15 and CAR15/TPC^KO^ NK was significantly improved compared to control NK treated animals (median survival in days: CAR15 vs Ctrl 63 vs 25, p<0.01; CAR15/TPC^KO^ vs Ctrl 30 vs 25, p<0.05); however, in contrast to the *in vitro* killing assays, we saw no significant differences between CAR15 NK and CAR15/TPC^KO^ NK (median survival in days: CAR15 vs CAR15/TPC^KO^ 63 vs 30, p=0.5982) **(Figure 6e)**. Among all 5 mice in the CAR15 group, four animals were euthanized due to paralysis and one animal due to rapid loss of body weight. Necropsy of the animal with rapid weight loss revealed a significantly enlarged ovary on the left side **(Supplement Figure 13c)**. For the CAR15/TPC^KO^ group, one animal survived long term (128 days) before presenting with paralysis, two animals were euthanized due to rapid body weight loss and were found to harbor enlarged ovaries and/or brain tumor masses. Two CAR15/TPC^KO^ treated animals were also euthanized due to rapid body weight loss with no significant tumor burden detected **(Figure 6b, Supplement Figure 13c)**. Histology of ovary and brain tumors indicated a domination of hCD19+ cells with no signs of hCD56+ NK cells **(Supplement Figure 13d)**.

**Fig. 6.**
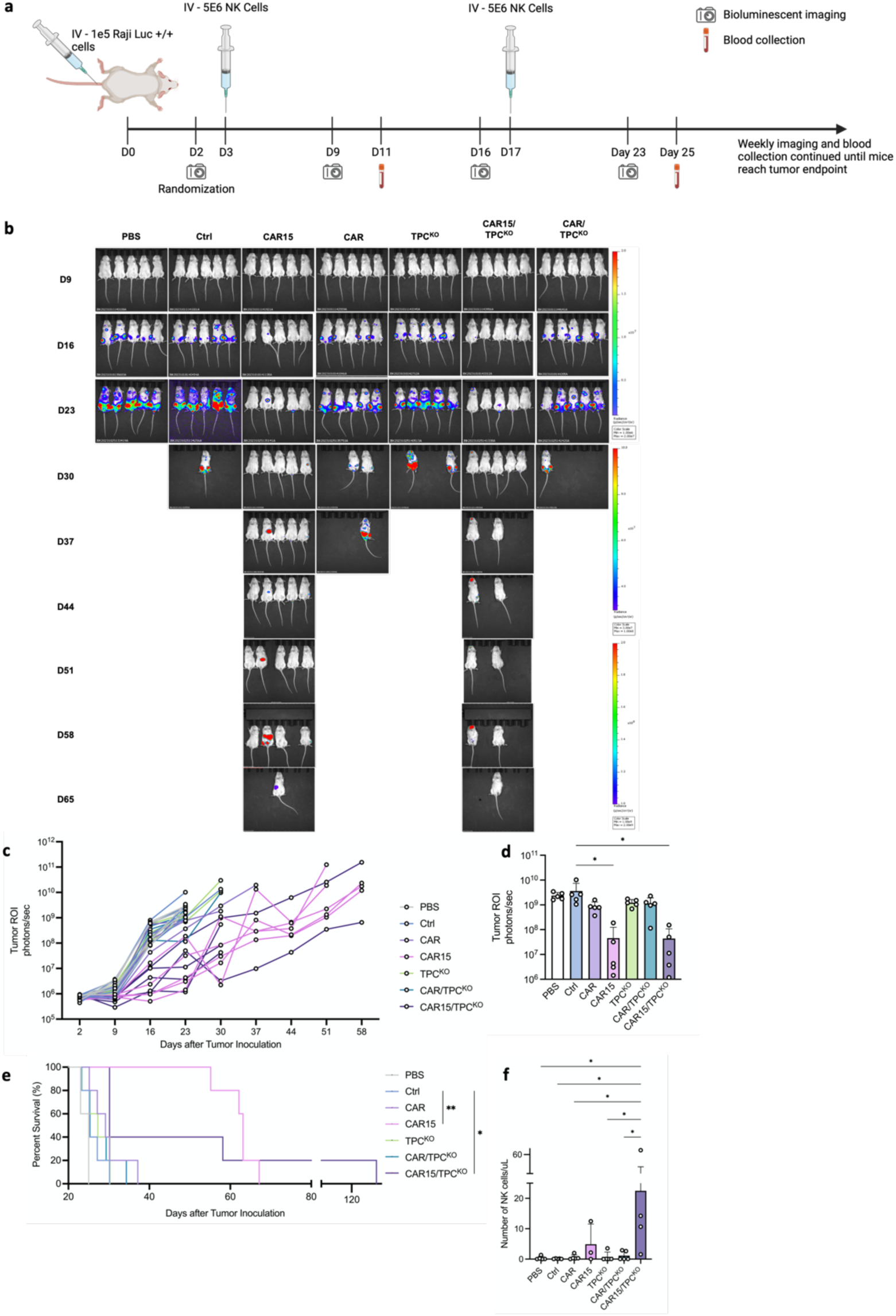
Multiplex edited CD19 CAR-NK cells are highly functional *in vivo.* **a** Schema of *in vivo* study design and timeline. **b-c** Luminance (ROI) of individual tumor burden of mice bearing Raji^hi/hi^ cells following treatment with PBS, Ctrl or engineered NK cells. **d** Tumor burden of each group on day 23, quantified by ROI (photons/sec) of each mouse (n=5). P-values calculated by one-way ANOVA test (*P ≤ 0.05). **e** Survival curve of each group shown in Kaplan-Meier curve (n=5). P-values calculated by Mantel-Cox test. Ctrl versus CAR15, **P ≤ 0.01; Ctrl versus CAR15/CTP^KO^, *P ≤ 0.05. **f** Numbers of circulating NK cells measured by NK cell number per uL of peripheral blood at end point. Data represented as mean ± SD.

Weekly peripheral blood (PB) collection and endpoint PB, bone marrow (BM) and spleen collections were performed to monitor the persistence of NK cells in each treatment group **(Figure 6f, Supplement Figure 13e, g&h)**. We observed a significantly higher level of NK cells in blood, BM and spleen at endpoint in CAR15/TPC^KO^ NK group than any other groups **(Figure 6f, Supplement Figure 13g, h)**. Interestingly, the CAR15 group had no detectable NK cells in the BM or spleen and minimal NK cells in PB. **(Figure 6f, Supplement Figure 12g, h)**. The amount of Raji^hi/hi^ cell in BM was also determined at endpoint, where we observed a significantly lower frequency of Raji^hi/hi^ cells in CAR15 NK and CAR15/TPC^KO^ NK groups compared to other groups, yet no significant difference was observed between the two groups **(Supplement Figure 12j)**. Together, we observed a significant improvement in tumor control and overall survival of mice treated with NK cells expressing sIL-15, with the CAR15/TPC^KO^ NK group exhibiting a slightly improved persistence but no survival advantages over CAR15 NK.

## Discussion

Cellular therapy using CAR-expressing primary effector cells has shown great success in the clinic and resulted in multiple FDA-approved treatments for hematologic malignancies and solid tumors^2^. These current-generation of FDA-approved CAR therapies largely utilize autologous 𝛼/𝛽 T cell chassis and viral vector engineering methods, which limits the scope of potential therapy recipients and results in high manufacturing costs and long timelines. A more appealing approach is to utilize a universal cell type, such as primary NK cells, that are engineered using non-viral methods. Further, as the landscape of cancer therapy shifts towards solid tumors, it is increasingly evident that methods such as cytokine armoring and immune checkpoint editing through genome engineering will be required to create more potent and persistent therapies. Since CRISPR/Cas9 was co-opted for genome editing over a decade ago, engineering efficiencies in primary immune cells have been significantly improved^81,82^. It remains, however, that indels and translocations caused by Cas9 induced DSBs could negatively impact cellular function and introduce major safety concerns for CRISPR/Cas9-based therapies^81,83^. In an effort to circumvent this issue, we applied base editing technology to NK cell engineering for the first time and demonstrated NK cells could be multiplex engineered with undetectable rates of insertions or deletions (indels). KO of our selected targets or installation of gain of function edits resulted in increased baseline functionality, which could be combined to further increase performance. We then combined multiplex base editing with non-viral transposon engineering to generate a CAR-NK engineering pipeline for precision enhancement of next generation CAR-NK cell products. These multiplex base-edited CAR-NK cells showed improved functionality *in vitro* and *in vivo* against a challenging model of Burkitt’s lymphoma model and provide compelling pre-clinical evidence for a future NK based CAR therapy option.

Our initial engineering studies examined whether NK cells were amenable to individual targeted base editing at high-frequency and how these edits impact NK cell function. We began with selecting a variety of targets that play inhibitory but critical roles in NK cell function with an emphasis on clinical relevance. To confirm that successful genetic modifications lead to protein loss and functional enhancements, we validated protein loss and functional improvement as part of the validation process, respectively. Other than *KLRG1*, all inhibitory target KOs exhibited the predicted functional enhancement. *KLRG1* KO NK cells’ failure to demonstrate improved function is in line with some previous studies disrupting *KLRG1* in NK cells^84^. As speculated in those studies, this contradictory finding may be attributed to a relatively low inhibitory potential of *KLRG1* on NK cell function^84,85^.

Our inability to detect PD-1 protein on the surface of NK cells is in line with numerous previous reports and reflects the ongoing controversy of the role of PD-1 in NK cells^20,50,86,87^. For instance, the Moretta group has published multiple studies on this issue and found that NK cells constantly produce PD-1 mRNA and protein, but only rapidly externalize PD-1 protein upon activation by certain stimuli^86^. Davis *et. al.* also reported minimal surface expression of PD-1 in NK cells from healthy individuals^87^. Moreover, several previous studies have used functional assays with anti-PD-1 agents to validate PD-1 expression on the NK cell surface, and our results here demonstrating significant cytotoxicity improvement of *PDCD1* KO NK cells against *PD-L1*+ target cells further support the role of PDCD1 in NK cell inhibition by PD-L1^20,50^.

We also demonstrate the ability to install gain-of-function mutations in NK cells using base editing. The rapid downregulation of CD16a as a result of a proteolytic cleavage process upon NK cell activation serves as a negative feedback mechanism for ADCC. In a tumor microenvironment (TME) setting, functional inactivation of tumor associated NK cells reflects a further down- regulation of CD16a that significantly compromises NK cell cytolytic function against tumor cells^88,89^. BE installation of a non-cleavable CD16a improved both ADCC mediated killing and cytokine production in NK cells. Notably, this BE mediated approach is likely significantly safer and more physiologically relevant compared to integration of a ncCD16A cDNA expression cassette using viral delivery^64^. Unexpectedly, we also observed a significant improvement in basal killing of Raji cells with ncCD16A NK cells. We speculate that the constitutive expression of the non-cleavable CD16a may result in low level tonic signaling, contributing to this phenomenon. This phenomenon could further benefit patients with compromised NK cytolytic functions, and a closer look at the downstream mechanisms of this phenomenon is warranted in future studies.

Moving forward to multiplex editing, we observed no significant reduction of editing efficiencies with up to six co-delivered sgRNAs. Surprisingly, there was an improvement of *KLRG1* KO efficiency at the multiplex level compared to the single-target experiments. One explanation for this phenomenon is a potential ‘carrier effect’ linked to increasing the total amount of co-delivered sgRNAs. This is similar to previous studies reporting a higher extracellular DNA concentration leads to a higher amount of DNA delivered to the nucleus^90,91^. However, further investigation is required to illuminate the exact mechanisms behind this observation.

To examine whether multiplex editing resulted in enhanced *in vitro* function, we developed and utilized an engineered Burkitt’s lymphoma cell line Raji^hi/hi^ as a proof-of-principle model. We hypothesized that the number of inhibitory gene KOs would positively correlate with the improvement of NK cell function, and selected *AHR*, *CISH*, *TIGIT* and *PDCD1* as candidates. *KLRG1* was excluded due to its failure to demonstrate functional improvement at single KO level. Contrary to our hypothesis, the TPC KO group demonstrated the most significant improvement in cytotoxicity against Raji^hi/hi^ cells. Additionally, the fact that no significant differences in killing were observed among TP, TPA, and TPAC KO groups throughout all E:T ratios is another interesting finding that warrants further investigation. It is possible that Raji^hi/hi^ cells produced lower amounts of AhR ligand (kynurenine) than K562, leading to a less pronounced inhibitory effect on NK cells. These findings suggest that the number of inhibitory gene KOs does not always correlate with improved NK cell functionality, and an optimization of the KO combination is likely required for precision enhancement against each unique tumor type of interest.

Finally, in an effort to test the impacts of multiplex editing on CAR-NK cell function, we co-delivered a CD19 CAR with multiplex base editing using the *TcBuster* DNA transposon system. Compared to traditional virus-based delivery methods, non-viral transposon systems significantly reduce the manufacturing cost and timelines. Integration efficiencies of CD19 CAR constructs were correlated with its cargo size, and are comparable to that of virus-based delivery methods^19,92^. One unexpected finding is the significantly higher CD19 CAR integration rate in the groups with TPC KO. We suspect a similar carrier effect may be contributing to the higher rates; however, further investigation is necessary to define the mechanism behind this observation^90,91^. Multiple previous studies have reported improved *in vivo* persistence of CAR-NK cells with sIL-15 expression^17–19,93^. Here, we adopted this idea in an attempt to improve proliferation and prolong the persistence of NK cell products *in vivo* by armoring cells with sIL-15. Our animal study clearly demonstrated a strong correlation of IL-15 expression with NK cell persistence *in vivo*, which translates into a significantly delayed tumor progression and improved overall survival compared to groups without IL-15 expression. For groups treated with NK cells expressing sIL-15, one animal from group CAR15 developed a tumor in the ovary, and two animals from group CAR15/TPC^KO^ developed a tumor in the ovary and/or brain. The fact that all these tumors developed in known immune privileged areas serves as indirect evidence of robust tumor clearance in peripheral blood and BM where full immune surveillance is expected^94^. The CAR15/TPC^KO^ group’s failure to exhibit an improved survival over CAR15 group is unexpected based on our *in vitro* results and thus requires closer examination. Among all 5 animals in the CAR15/TPC^KO^ group, two died with relatively low tumor burden but met endpoint criteria of significant decrease of body weight. We initially suspected cytokine release syndrome (CRS) in these mice; however, there was no significant increase in CRS related cytokines in serum at the endpoint (data not shown). A closer look at the weekly NK cell density data revealed a rapid NK cell proliferation after day 25 in 4 out 5 mice in CAR15/TPC^KO^ group, and 3 of those mice died in the following week. We suspect that this dramatic NK cell proliferation, not the absolute number of NK cells, led to the observed toxicity and eventual death in those mice, as the absolute number of NK cells in CAR15 group was equal to, if not higher, than that of the CAR15/TPC^KO^ group. Our observation is consistent with a previous study using sIL-15 expressing CAR-NK cells, which also found negative evidence of CRS and attributed the observed systemic toxicity to rapid NK cell proliferation *in vivo*^93^. It is worth noting that the only mouse with full tumor clearance was treated with CAR15/TPC^KO^ NK cells, suggesting a potential benefit of CAR15/TPC^KO^ over CAR15 treatment. However, further evaluation of the CAR15/TPC^KO^ NK treatment will be required to confirm improved tumor clearance and survival benefit over CAR15 NK treatment.

In summary, we reported the first application of BE in primary NK cells, with high editing efficiency for both single and multiplex editing. Moreover, with co-delivery of a mult-cistronic CAR construct using a non-viral transposon system, we present a highly flexible, fully non-viral engineering platform that allows precision enhancement of NK-based cell therapy against various types of cancer. With our proof-of-principle model, we have generated a pipeline for developing a highly customized NK product for any specific type of cancer. Moving forward, we are actively exploring the application of this platform into the frontier of solid tumors, including enhancement of NK cell homing to tumor sites, NK cell resistance to oxidative stress, and more.

## Methods

### Production of editing reagents

ABE8e plasmid was obtained from Addgene (https://www.addgene.org/138489/) and cloned into a pmRNA vector. ABE8e mRNA was produced by Trilink Biotechnologies. Nanoplasmids for CD19 CAR were synthesized by Aldevron.

### Guide RNA design

Guide RNAs (sgRNAs) for KO were designed using the base editing splice-site disruption sgRNA design program SpliceR (https://z.umn.edu/splicer). SpliceR is written in the R statistical programming language (v. 3.4.3). Briefly, SpliceR takes a target Ensembl transcript ID, a base editor PAM variant, and a species as an input. Using the exon and intron sequences from Ensembl, the program extracts the region surrounding every splice site based on a user-specified window. The pattern of N20-NGG is then matched to the antisense strand of the extracted sequence. Matched patterns are then scored based on the position of the target motif within the predicted editing window based on previous publications. Subsequently, sgRNAs are scored based on their position within the transcript, where sgRNAs earlier in the transcript receive a higher score. sgRNA for ncCD16a was designed manually to achieve the S197P mutation.

### Donor NK cell isolation

Peripheral blood mononuclear cells (PBMCs) from healthy human donors were obtained by automated leukapheresis (Memorial Blood Centers, Minneapolis, MN). CD56+CD3- NK cells were isolated from the PBMC population using the EasySep Human NK Cell Isolation Kit (STEMCELL Technologies, Cambridge, MA). NK cells were frozen at 2.5e6 or 5E6 cells/mL in CryoStor CS10 (STEMCELL Technologies, Cambridge, MA).

### NK cell culture

NK cells were cultured in CTS AIM VTM (ThermoFisher, Waltham, MA) with 5% CTS Immune cell SR (ThermoFisher, Waltham, MA), and 100 IU/mL IL-2 (PeproTech, Rocky Hill, NJ). NK cells were activated by co-culture with X-irradiated (100 Gray) feeder cells (K562 expressing membrane-bound IL-21 and 41BB-L) at various feeder to NK ratios (2:1 prior to electroporation, 5:1 72 hours after Neon electroporation, or 1:1 immediately after MaxCyte electroporation).

### NK cell electroporation

#### Neon electroporation

Feeder cell-activated NK cells were washed once with PBS (Ca^2+^ and Mg^2+^ free) and resuspended at 3E7 cells/mL in electroporation buffer. Protector RNase inhibitor (Sigma Aldrich, St. Louis, MO) was added to the mixture at a concentration of 0.8 U/µL and incubated for 5 minutes at room temperature. The cell mixture was added to 1.5ug of ABE8e mRNA and 1 nmol sgRNA on ice. This mixture was electroporated in a 10 µL tip using the Neon Transfection System (ThermoFisher, Waltham, MA) under the following conditions: 1825 volts, pulse width of 10 ms, 2 pulses. NK cells were allowed to recover at a density of 1E6 cells/mL in antibiotic-free medium containing 100 IU/mL IL-2 (PeproTech, Rocky Hill, NJ) for 72 hours at 37°C, before expansion with feeder cells at a 5:1 feeder to NK ratio.

#### MaxCyte electroporation

NK cells reaching 10-fold increase in cell number after feeder cell-expansion were washed once with electroporation buffer and resuspended at 1.75E8-1.00E9 cells/mL. Protector RNase inhibitor (Sigma Aldrich, St. Louis, MO) was added to the mixture at a concentration of 0.8 U/µL and incubated for 5 minutes at room temperature. The cell mixture was added to 4 ug of ABE8e mRNA, 5ug sgRNA, 3 µg of transposase mRNA and 5 µg transposon nanoplasmid on ice. This mixture was electroporated in a R-50x3 assembly using the MaxCyte System (ThermoFisher, Waltham, MA) under Expanded T Cell 4 protocol. NK cells were allowed to recover at a density of 1E6 cells/mL in antibiotic-free medium for 1 hour at 37°C, before expansion with feeder cells at a 1:1 feeder to NK ratio.

### Genomic DNA analysis

NK cells were harvested 10 days after electroporation, followed by DNA isolation and PCR amplification of CRISPR-targeted loci using Phire Tissue Direct PCR Master Mix kit (ThermoFisher, Waltham, MA). Base editing efficiency was analyzed at the genomic level by Sanger sequencing of the PCR amplicons, and subsequent analysis of the Sanger sequencing traces using the web app EditR (baseeditr.com)^41^.

### Primer and sequence

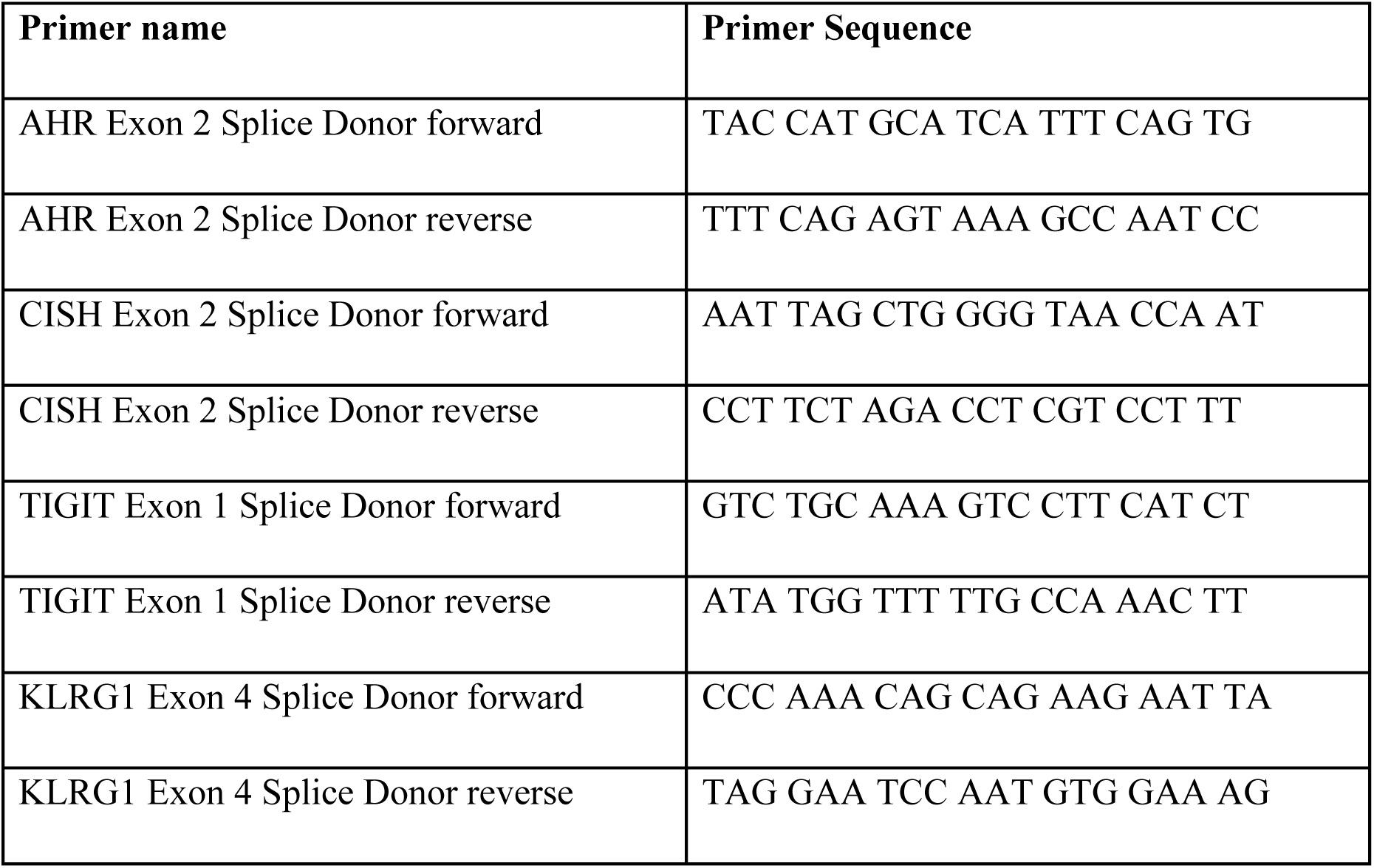
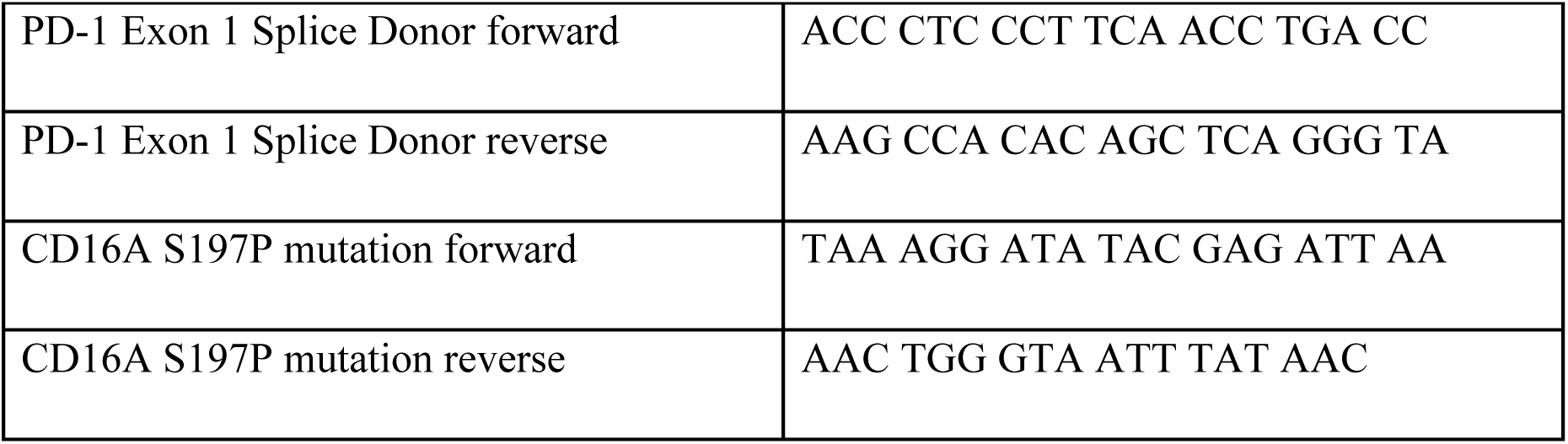

### Antibodies and flow cytometry

The following antibodies and dyes were used: APC-, FITC-, or BV650-conjugated anti-CD56 (clone 5.1H11, BioLegend; clone REA196, Miltenyi Biotec; or clone HCD56, BioLegend), FITC-conjugated anti-CD3 (clone OKT3; BioLegend), PE/Cy7-conjugated anti-PD-1 (clone EH12.2H7; BioLegend), eFluor 450- or Alexa FluorTM 700-conjugated anti-TIGIT (clone MBSA43; eBioscience), PE/DazzleTM 594- or FITC-conjugated anti-human CD45 (clone 14C2A07, BioLegend; clone SA231A2, BioLegend), PE/Cy7 anti-human CD16 (clone 3G8, BioLegend), BV421-conjugated anti-human CD45 (clone 2D1; BioLegend), BV605-conjugated anti-mouse CD45 (clone 30-F11; BioLegend), Brilliant violet 421-conjugated anti-IFNγ (clone 4S.B3; BioLegend), Alexa FluorTM 700-conjugated anti-TNF-⍺ (clone MAb11; BioLegend), PE-conjugated anti-CD34 (clone QBEnd-10, Invitrogen), APC-conjugated anti-PD-L1 (clone B7-H1, BioLegend), FITC-conjugated anti-CD155 (clone SKII.4, BioLegend), FITC-conjugated anti-DNAM-1 (clone 11A8, BioLegend), Brilliant violet 510-conjugated anti-CD161 (clone HP-3G10, BioLegend), PE/Cyanine7-conjugated anti-Tim3 (clone F38-2E2, BioLegend), APC-conjugated anti-NKG2D (clone 1D11, BioLegend), PE-conjugated anti-NKG2A (clone S19004C, BioLegend), Alexa FluorTM 700-conjugated anti-NKp46 (clone 9E2, BioLegend), PE/DazzleTM 594-conjugated anti-NKp30 (clone P30-15, BioLegend), FITC-conjugated anti-CD107a (clone H4A3; BD Biosciences), SYTOX Blue dead cell stain (ThermoFisher), Fixable viability dye eFluor 780 (eBioscience). Flow cytometry were performed on a CytoFLEX S flow cytometer (Beckman Coulter) and all data were analyzed using FlowJo version 10.4 software (FlowJo LLC).

### Immunoblotting assay

Proteins were isolated from 1E6 cells in complete RIPA buffer with protease and phosphatase inhibitors (Sigma-Aldrich, St. Louis, MO). Total protein was quantified using the Pierce BCA Protein Assay Kit (ThermoFisher, Waltham, MA) according to the manufacturer’s protocol. 3μg/μL of cell lysate was run and analyzed on the Wes platform after being denatured at 95 °C for 5 min according to the manufacturer’s protocol (ProteinSimple, San Jose, CA). Primary antibodies against CISH (Cell Signaling #8731), AhR (Cell Signaling #83200) and beta-actin (Cell Signaling #3700) were all used at 1:50 dilutions in kit-supplied buffer and platform-optimized secondary antibodies were purchased from ProteinSimple.

### NK cell intracellular cytokine staining (ICS) assay

Activated NK cells were plated at 1E6 cells/mL in NK cell medium without cytokines. After incubation overnight, the Burkitt’s Lymphoma cell line Raji or T-cell leukemia cell line Jurkat was added at an effector-to-target (E:T) ratio of 1:1. FITC-conjugated anti-CD107a was added to the culture and cells were incubated for 1 hour at 37°C. Brefeldin A and monensin (BD Biosciences, San Jose, CA) were added and cells were incubated for 4 hours. Cells were stained with fixable viability dye, then for extracellular antigens. Cells were fixed and permeabilized using BD Cytofix/Cytoperm (BD Biosciences, San Jose, CA) following manufacturer’s instructions. Cells were then stained for intracellular IFN-γ and/or TNF-⍺ and analyzed by flow cytometry.

### NK cell cytotoxicity assays

Cancer cell lines were seeded into a black round-bottom 96-well plate (5E4 cells per well). NK cells were added to the wells in triplicate at the indicated E:T ratios. Target cells without effectors served as a negative control (spontaneous cell death) and target cells incubated with 1% Triton X- 100 served as a positive control (maximum killing). Co-cultures were incubated at 37°C for 48 hours. After incubation, D-luciferin (potassium salt; Gold Biotechnology, St. Louis, MO) was added to each well at a final concentration of 25 ug/mL and incubated for 5 minutes with gentle shaking. Luminescence was read in endpoint mode using a BioTek Synergy microplate reader. For *in vitro* killing assays, percent CAR expression was equalized across groups using matched NK cells (for CD19 CAR only groups) or TPC^KO^ NK cells (for co-delivery groups), before co-culture set up.

### Antibody-dependent cellular cytotoxicity assay

CD20+ Raji cells were pre-treated with anti-hCD20 mAb (InvivoGen, San Diego, CA) at 10 μg/mL for 30 min at 37°C, washed with PBS, and seeded into a black round-bottom 96-well plate (5E4 cells per well). NK cells were added to the wells in triplicate at the indicated E:T ratios. Co-cultures were incubated at 37°C for 4 hours. After incubation, D-luciferin (potassium salt; Gold Biotechnology, St. Louis, MO) was added to each well at a final concentration of 25 ug/mL and incubated for 5 minutes with gentle shaking. Luminescence was read in endpoint mode using a BioTek Synergy microplate reader.

### Transposon integration sequencing

Genomic DNA was isolated from engineered CAR-NK cells as previously described^95^. Briefly, sequencing libraries were prepared from 150 ng genomic DNA quantified by Picogreen (Life Technologies) using the Lotus DNA Library Prep Kit (Integrated DNA Technologies) according to the manufacturer’s specifications for libraries undergoing target enrichment. Ligations used vendor-supplied “stubby” adapters, with sample-specific 8-bp unique dual indices (UDIs) added during final library amplification. Hybridization capture was performed per manufacturer’s protocol with up to 16 libraries in multiplex (500ng per library) using xGen universal blocking oligos (IDT) and a custom biotinylated xGen oligo probe pool designed to hybridize to the inserted transposon sequence. Given the small probe panel size, hybridizations and temperature-sensitive washes were performed at 63 °C and the total hybridization time was increased from 4 h to 16 h. Captured libraries were then amplified to ≥2 nM using KAPA HiFi HotStart 2X PCR master mix, quantified by Picogreen, sized on an Agilent TapeStation using the D1000 assay, normalized, and pooled for 150-bp paired-end sequencing on an Illumina NovaSeq* SP flowcell.

### Transposon integration analysis

We analyzed integration site data for TcBuster and compared results to published literature data for integration sites of other transposase and viral systems as previously described^95^. Specifically, comparison sequencing datasets were generated by outside sources using different experimental methods. Raw reads from comparison datasets were retrieved (Accession IDs: Lentivirus: SAMN11351981, SAMN11351982, SAMN11351983, SAMN11351984, SAMN00188192, SAMN00188193; Sleeping Beauty: SAMN02870102; and PiggyBac: SAMN02870101) and computationally mapped and analyzed in the same manner as the in-house generated data using Python (v3.7.10, run on CentOS 7 Linux).

### *In vivo* study design

Specific pathogen-free female NOD-scid IL2Rgammanull (NSG) mice were purchased from The Jackson Laboratory (RRID:IMSR_JAX:005557). Tumor challenge studies were performed using the Raji-luc CD155hi/PD-L1hi cell line. Specifically, mice were implanted with 1E5 Raji cells resuspended in PBS and delivered in 100 µL via tail vein. Two days post tumor implantation, the mice were randomized into treatment groups (n=5) and received either PBS, non-engineered (control) NK cells, CAR NK cells, CAR15 NK cells, CTP^KO^ NK cells, CAR/CTP^KO^ NK cells, or CAR15/CTP^KO^ NK cells the next day. For each group, all tumor growth was monitored by weekly bioluminescence imaging of mice 5 minutes post IP injection of D-luciferin (100 µL total volume, 28 mg/kg) using an IVIS® Spectrum *in vivo* imaging system followed by ROI analysis of tumor images (Living Image software, version 4.7.3). NK cell persistence in peripheral blood was monitored by weekly submandibular blood collection. Endpoint criteria are either mouse paralysis or >20% weight loss within a week. At endpoint, peripheral blood, spleen, and bone marrow were collected and processed to single cell suspensions through standard mashing and ACK-processing and stained for phenotyping markers. This study was carried out in strict accordance with the recommendations in the Guide for the Care and Use of Laboratory Animals of the National Institutes of Health. The protocol and all procedures were approved by the University of Minnesota Institutional Animal Care and Use Committee (Protocol #2110-39527A). The health of the mice was monitored daily by University of Minnesota veterinary staff.

### Statistical analysis

The Student’s t-test was used to evaluate the significance of differences between the two groups. Differences between three or more groups with one data point were evaluated by a one-way ANOVA test. Differences between three or more groups with multiple data points were evaluated by a two-way ANOVA test. Differences between groups in our *in vivo* study were evaluated by the Log-rank (Mantel-Cox) test. The level of significance was set at α = 0.05. All statistical analyses were performed using GraphPad Prism 9.2.0.

## Acknowledgements

B.S.M. is supported by the following NIH grants (R01AI146009, R01AI161017, P01CA254849, P50CA136393, U24OD026641, U54CA232561, P30CA077598, U54CA268069), and received support from Children’s Cancer Research Fund, the Fanconi Anemia Research Fund, and the Randy Shaver Cancer Research and Community Fund. B.R.W. is supported by the following NIH grants (R21CA237789, R21AI163731, P01CA254849, P50CA136393, U54CA268069, R01AI146009), and received support from Alex’s Lemonade Stand Foundation, Children’s Cancer Research Fund, and Rein in Sarcoma.

