## Supplement Figures for "Precision Enhancement of CAR-NK Cells through Non-Viral Engineering and Highly Multiplexed Base Editing"

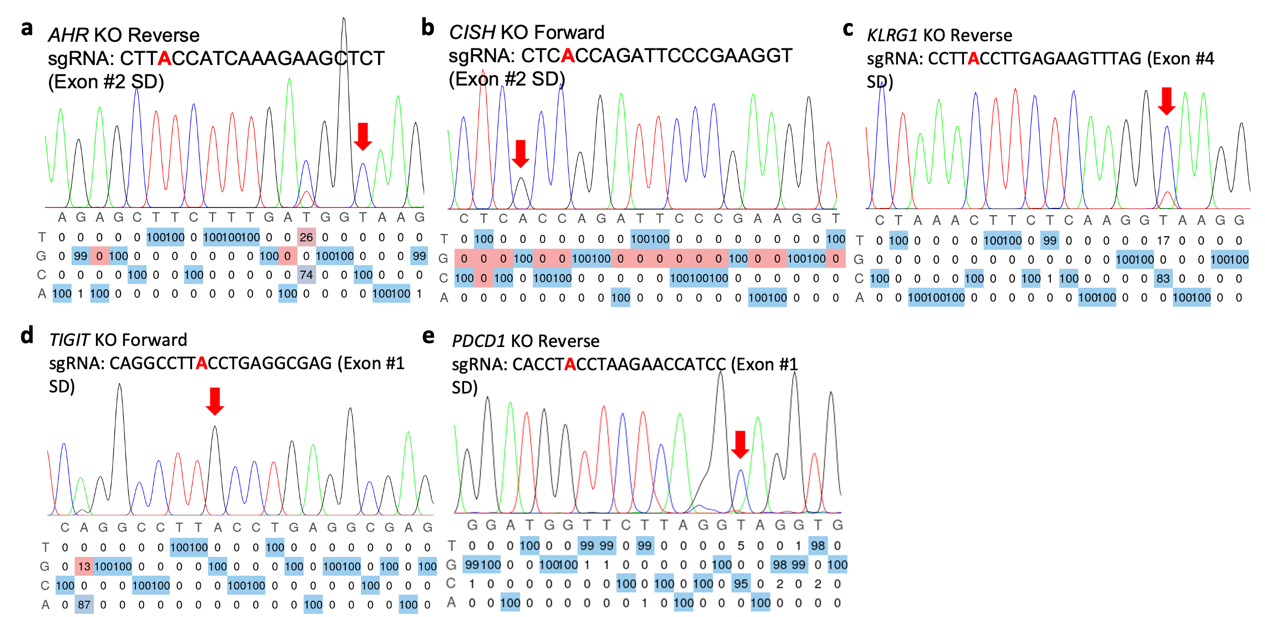

**Sup Fig. 1 Representing Sanger sequencing result of each single edit in NK cells using BE.** **a** Representing Sanger sequencing result of *AHR* KO showing 100% A to G conversion. **b** Representing Sanger sequencing result of *CISH* KO showing 100% A to G conversion. **c** Representing Sanger sequencing result of *KLRG1* KO showing 83% A to G conversion. **d** Representing Sanger sequencing result of *TIGIT* KO showing 100% A to G conversion. **e** Representing Sanger sequencing result of *PDCD1* KO showing 100% A to G conversion.

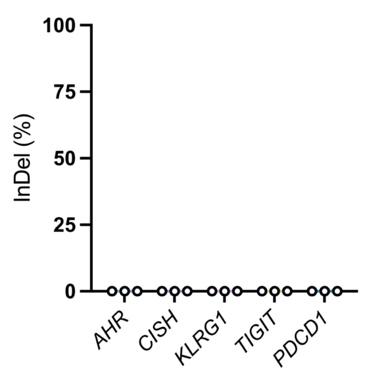

**Sup Fig. 2 Zero indels in single gene KO in NK cells using BE.** Indel analysis was quantified using Inference of CRISPR Edits (ICE).

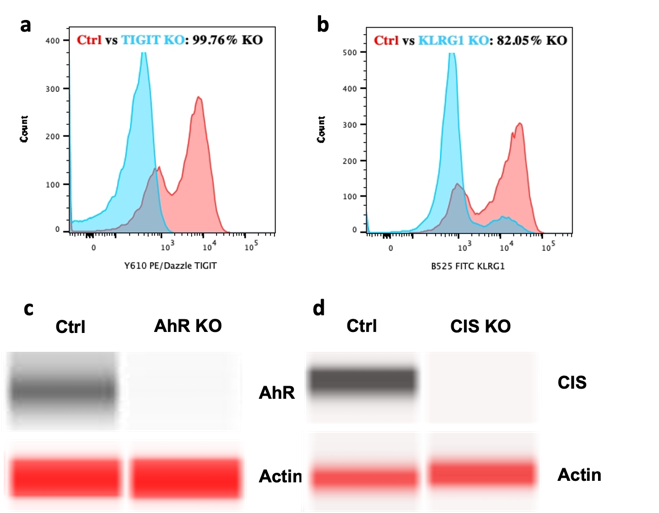

**Sup Fig. 3 Representing protein level result of each single edit in NK cells using BE. a** Representative flow plot showing protein level editing efficiency of *TIGIT* KO (Ctrl in red and KO in blue). **b** Representative flow plot showing protein level editing efficiency of *KLRG1* KO (Ctrl in red and KO in blue). **c** Representative western blot result showing protein level editing efficiency of *AHR* KO. **d** Representative western blot result showing protein level editing efficiency of *CISH* KO.

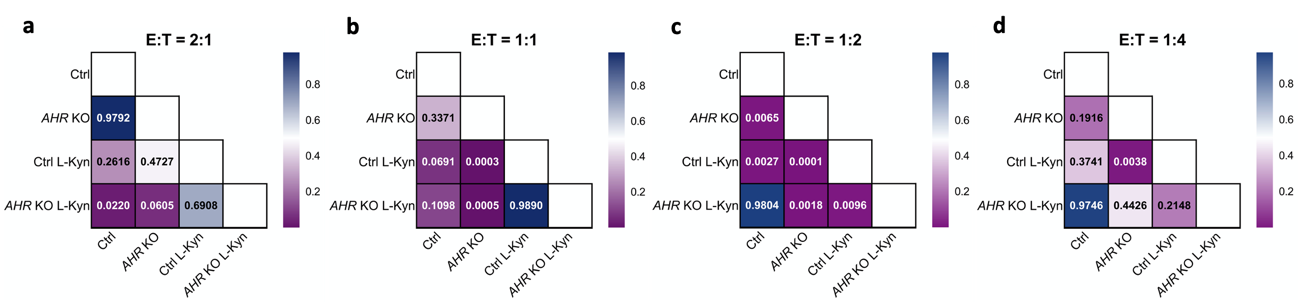

**Sup Fig. 4 Statistical significance of different E:T ratios for AHR single KO functional assay. a** Statistical significance (P-value) between each condition of AHR KO functional killing assay at E to T ratio of 2:1. **b** Statistical significance (P-value) between each condition of AHR KO functional killing assay at E to T ratio of 1:1. **c** Statistical significance (P-value) between each condition of AHR KO functional killing assay at E to T ratio of 1:2. **d** Statistical significance (P-value) between each condition of AHR KO functional killing assay at E to T ratio of 1:4.

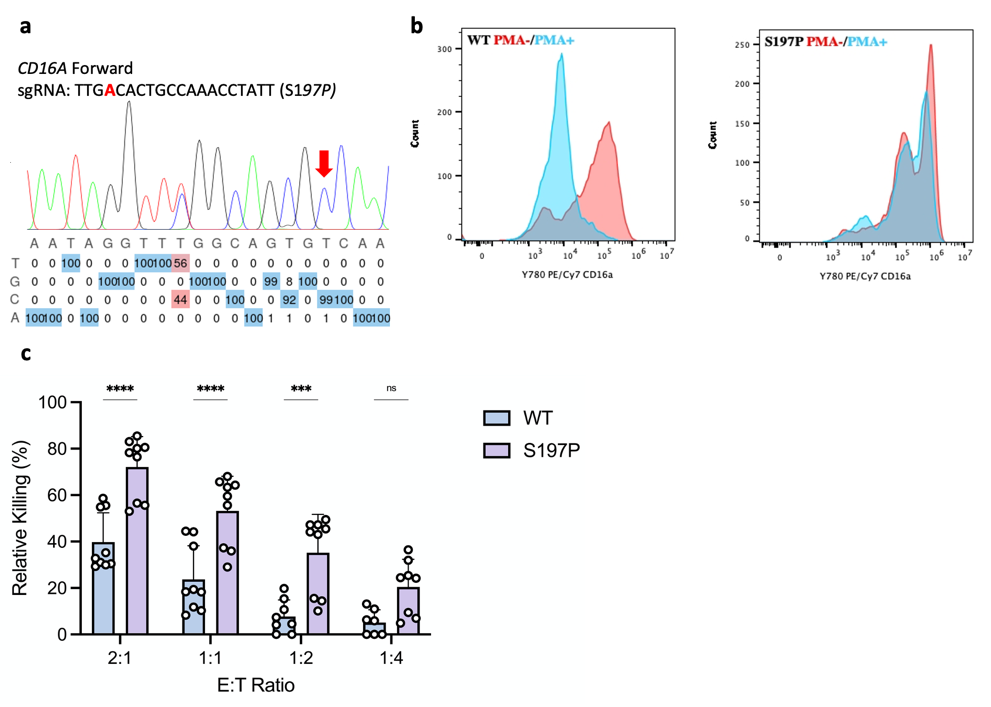

**Sup Fig. 5 Additional ICS and killing ADCC assay results of ncCD16a NK cells. a** Representing Sanger sequencing result of non-cleavable CD16A showing 100% A to G conversion. **b** Representative flow plots showing CD16a retention when treated with or without PMA (without PMA in red and with PMA in blue). **c** Ability of S197P CD16a versus WT CD16a NK cells to kill CD20+ Raji cells at various E:T ratios as measured by luciferase luminescence assay. Assays run in triplicate in n=3 independent biological NK cell donors. Data represented as mean ± SD. P-values calculated by two-way ANOVA test (***P ≤ 0.001, ****P ≤ 0.0001)

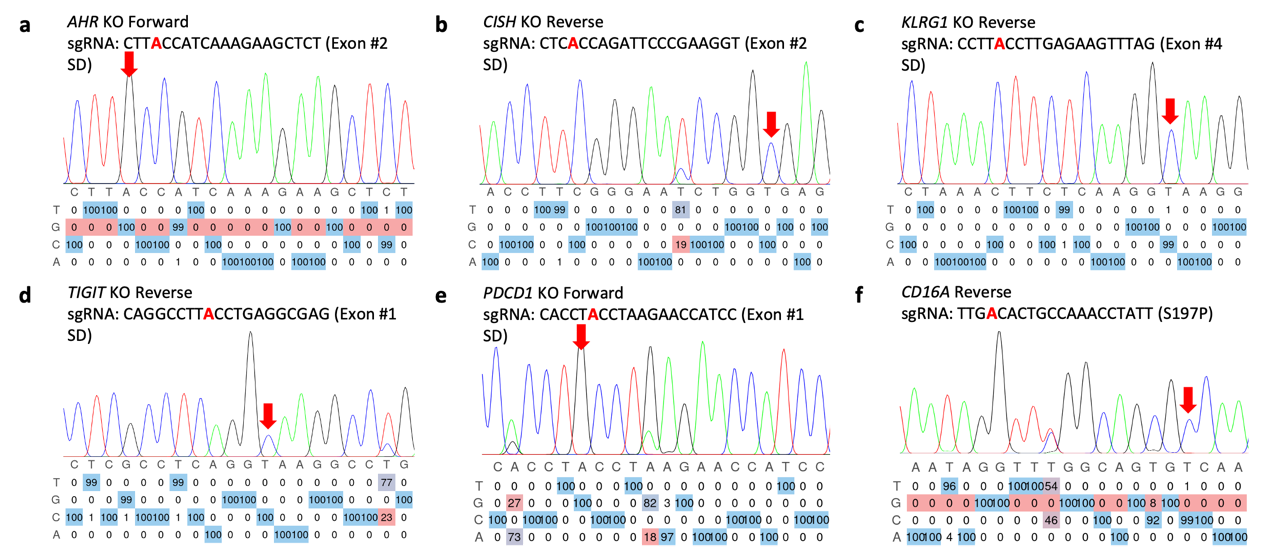

**Sup Fig. 6 Representing Sanger sequencing result of multiplex edit in NK cells using BE.** **a** Representing Sanger sequencing result of *AHR* KO showing 100% A to G conversion for X6 KO condition. **b** Representing Sanger sequencing result of *CISH* KO showing 100% A to G conversion for X6 KO condition. **c** Representing Sanger sequencing result of *KLRG1* KO showing 99% A to G conversion for X6 KO condition. **d** Representing Sanger sequencing result of *TIGIT* KO showing 100% A to G conversion for X6 KO condition. **e** Representing Sanger sequencing result of *PDCD1* KO showing 100% A to G conversion for X6 KO condition. **f** Representing Sanger sequencing result of non-cleavable *CD16A* showing 99% A to G conversion for X6 KO condition.

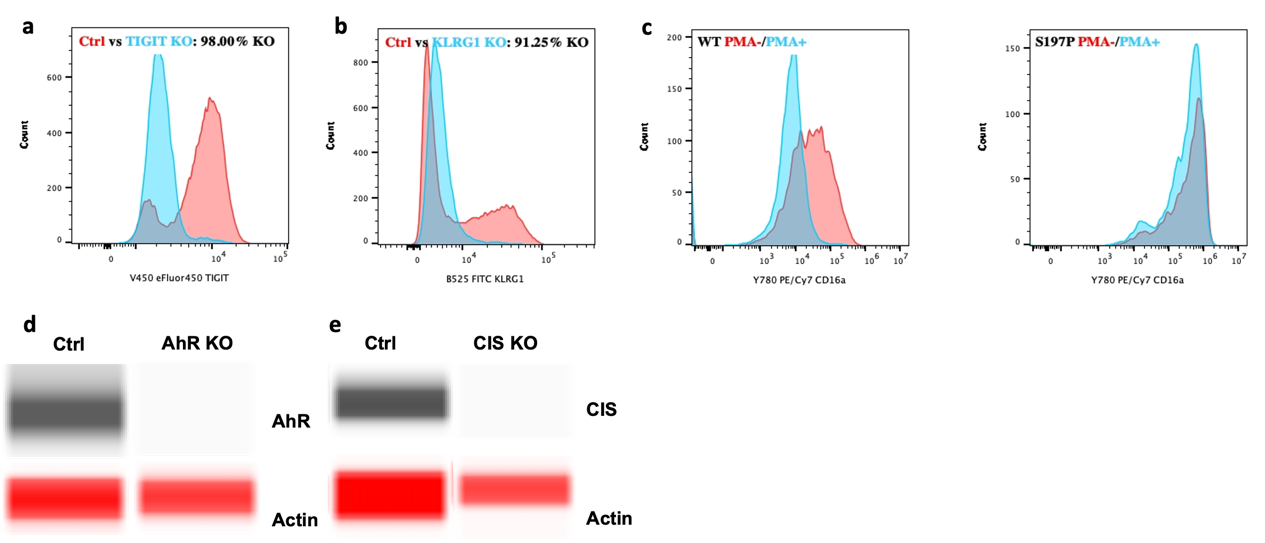

**Sup Fig. 7 Representing protein level result of multiplex edit in NK cells using BE. a** Representative flow plot showing protein level editing efficiency of *TIGIT* KO (Ctrl in red and KO in blue) for X6 KO condition. **b** Representative flow plot showing protein level editing efficiency of *KLRG1* KO (Ctrl in red and KO in blue) for X6 KO condition. **c** Representative flow plots showing CD16a retention when treated with or without PMA (without PMA in red and with PMA in blue) for X6 KO condition. **d** Representative digital western blot result showing protein level editing efficiency of *AHR* KO for X6 KO condition. **e** Representative digital western blot result showing protein level editing efficiency of *CISH* KO for X6 KO condition.

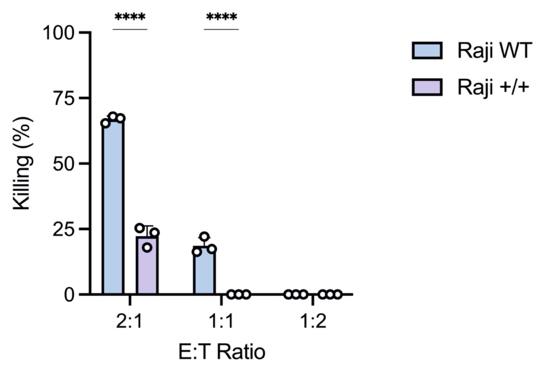

**Sup Fig. 8 Raji^hi/hi^ cell line exhibits resistance of NK killing than WT Raji cell line.** Ability of non-engineered NK cells to Raji WT or Raji^hi/hi^ cells at various E:T ratios as measured by luciferase luminescence assay. Assays run in triplicate in n=1 independent biological NK cell donors. Data represented as mean ± SD. P-values calculated by one-way ANOVA test (****P ≤ 0.0001).

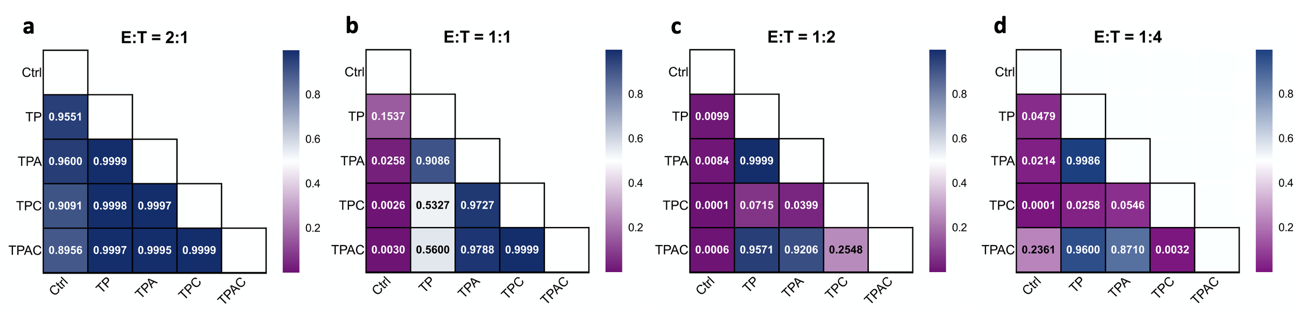

**Sup Fig. 9 Statistical significance of different E:T ratios for multiplex optimization killing assay. a** Functional killing assay statistical significance (P-value) between each KO combination at E to T ratio of 2:1. **b** Functional killing assay statistical significance (P-value) between each KO combination at E to T ratio of 1:1. **c** Functional killing assay statistical significance (P-value) between each KO combination at E to T ratio of 1:2. **d** Functional killing assay statistical significance (P-value) between each KO combination at E to T ratio of 1:4.

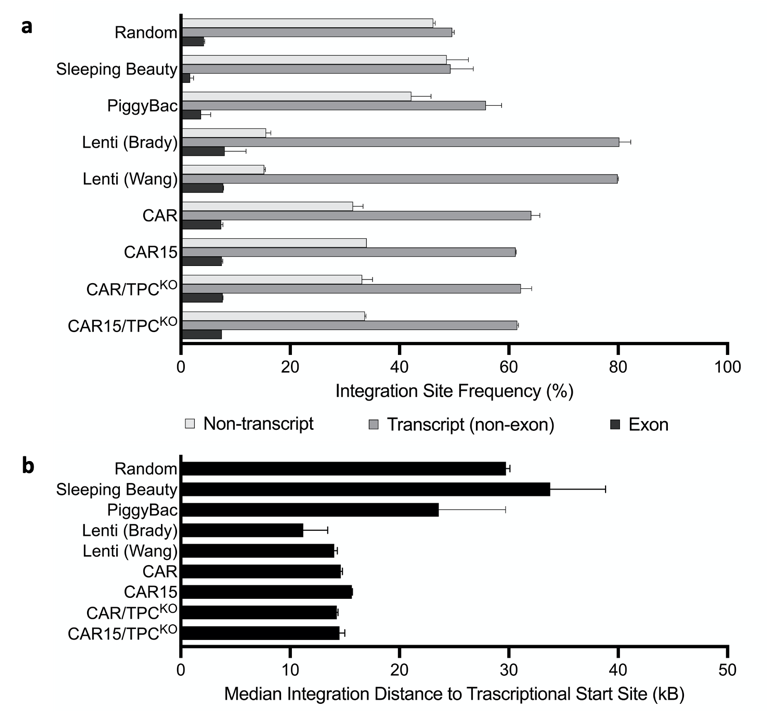

**Sup Fig. 10 Integration site analysis reveals reduced transcript integration** **with TcB. a** Analysis of the frequency of integration events in exon-coding transcript sites, non-exon-coding transcript sites, and non-transcript sites. Samples were collected from primary human NK cells transfected with TcBuster (TcB) or historical data sets generated by Wang et al., Brady et al., and Gogol-Döring et al. Random in silico control sets were also generated and analyzed. Lenti: lentiviral dataset. Graph displays mean ± SD, biologically independent samples. n = 2 for CAR, CAR15, CAR/CTP^KO^, CAR15/CTP^KO^, and Lenti (Wang); n = 9 for Sleeping Beauty; n =9 for PiggyBac; n = 4 for LV (Brady); n = 12 for Random. **b** Median distance measured between transposon integration sites and the transcriptional start site of the nearest gene is presented for transposon and lentiviral integration. Lenti: lentiviral dataset. Graph displays mean ± SD, biologically independent samples. n = 2 for CAR, CAR15, CAR/CTP^KO^, CAR15/CTP^KO^, and Lenti (Wang); n = 9 for Sleeping Beauty; n =9 for PiggyBac; n = 4 for LV (Brady); n = 12 for Random.

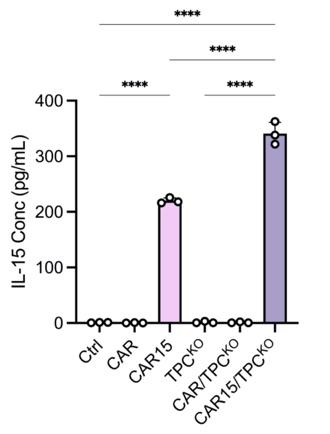

**Sup Fig. 11 ELISA results showing significantly higher sIL-15 expression in CD19 CAR IL-15 expression groups.** Assays run in triplicate in n=1 independent biological NK cell donors. Data represented as mean ± SD. P-values calculated by one-way ANOVA test (****P ≤ 0.0001).

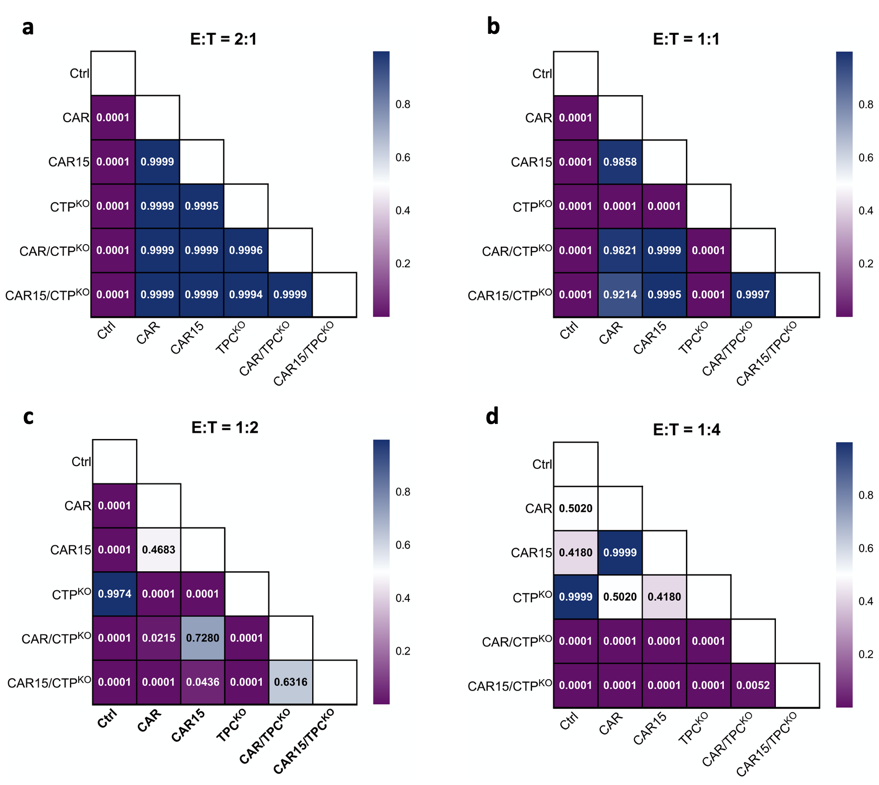

**Sup Fig. 12 Statistical significance of different E:T ratios for simultaneous BE and transposon CD19 CAR engineering killing assay. a** Killing assay statistical significance (P-value) between each condition at E to T ratio of 2:1. **b** Killing assay statistical significance (P-value) between each condition at E to T ratio of 1:1. **c** Killing assay statistical significance (P-value) between each condition at E to T ratio of 1:2. **d** Killing assay statistical significance (P-value) between each condition at E to T ratio of 1:4.

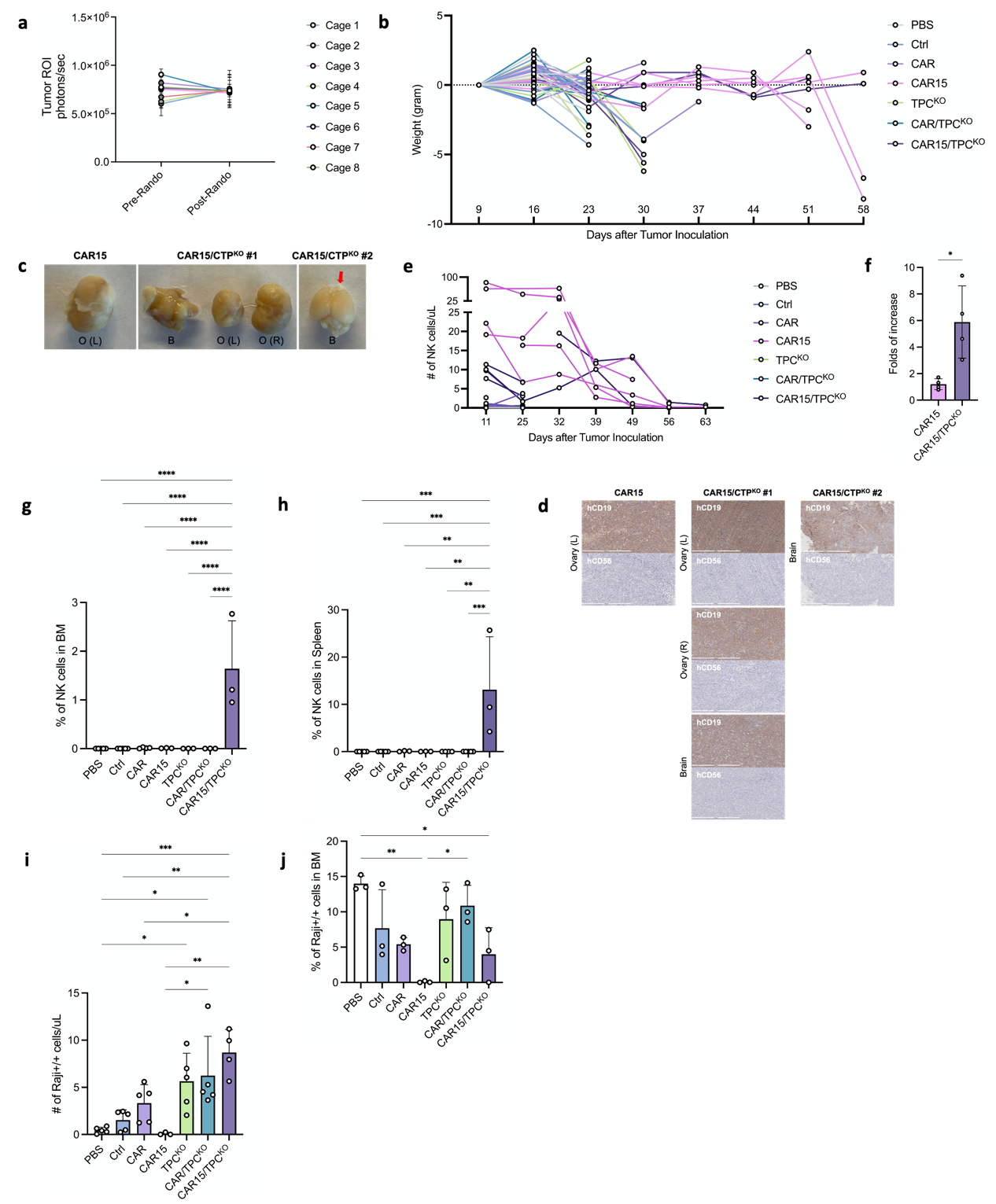

**Sup Fig. 13 Additional results supporting multiplex edited CD19 CAR-NK cells are highly functional *in vivo.* a** Pre-treatment randomization results showing similar average tumor burden (tumor ROI) were assigned to each treatment group. **b** Weekly body weight change of individual mice over time. **c** Pictures of ovarian and brain tumors found in group CAR15 and CAR15/TPC^KO^. O: Ovary, B: Brain; L: Left, R: Right. **d** Histology of ovarian and brain tumors found in group CAR15 and CAR15/TPCKO. Tumor slices of indicated groups were subjected to immunohistochemical analysis using anti-human CD19 and anti-human CD56. **e** Weekly monitoring of NK cells in peripheral blood of individual mice over time. **f** Quantification of NK cell proliferation of CAR15 and CAR15/TPC^KO^ groups between day 25 and 32 (folds of increase: day 32 NK cell count versus day 25 NK cell count). **g** Quantification of NK cells in BM at endpoint measured by percentage of NK cells in BM. **h** Quantification of NK cells in spleen at endpoint measured by percentage of NK cells in spleen. **i** Quantification of Raji^hi/hi^ cells in peripheral blood at endpoint measured by Raji^hi/hi^ cell count per uL of blood. **j** Quantification of Raji^hi/hi^ cells in BM at endpoint measured percentage of Raji^hi/hi^ in BM. *In vivo* study run in n=5 per group. Data represented as mean ± SD. P-values calculated by one-way ANOVA test (*P ≤ 0.05, **P ≤ 0.01, ***P ≤ 0.001, ****P ≤ 0.0001)
